# Golgi localized Arl15 regulates cargo transport, cell adhesion and motility

**DOI:** 10.1101/2022.08.18.504432

**Authors:** Prerna Sharma, Pooja Hoovina Venkatesh, Neha Paddillaya, Nikita Shah, BR Rajeshwari, Archishman Dakua, Aravind Penmatsa, Nagaraj Balasubramanian, Namrata Gundiah, Subba Rao Gangi Setty

## Abstract

Arf-like GTPases (Arls) regulate membrane trafficking and cytoskeletal organization. Genetic studies predicted a role for Arl15 in type-2 diabetes, insulin resistance, adiposity, and rheumatoid arthritis. Recent studies indicate a possible role for Arl15 in multiple physiological processes, including magnesium homeostasis. However, the molecular function of Arl15 is poorly defined. We evaluated the role of Arl15 in vesicular transport using techniques to quantify cargo trafficking, to mechanobiology. Fluorescence microscopy of stably expressing Arl15-GFP HeLa cells showed its localization to the Golgi and cell surface, including filopodia, and a cohort to recycling endosomes. The dissociation of Golgi, using small molecular inhibitors or the expression of Arf1 dominant-negative mutant, completely mislocalized Arl15 to the cytosol. Interestingly, site-directed mutagenesis analysis identified a novel V80A mutation in the GTP-binding domain that turns Arl15 into a dominant-negative form with reduced number of filopodia. Depletion of Arl15 in HeLa cells caused mislocalization of cargo, such as caveolin-2 and STX6, from the Golgi. Arl15 knockdown cells displayed reduced filopodial number, altered focal adhesion kinase organization, and enhanced soluble and receptor-mediated cargo uptake without affecting the TfR recycling. Arl15 knockdown decreased cell migration and enhanced cell spreading and adhesion strength. Traction force microscopy experiments revealed that Arl15 depleted cells exert higher tractions and generate multiple focal adhesion points during the initial phase of cell adhesion as compared to control cells. Collectively, these studies demonstrated a functional role for Arl15 in the Golgi, which includes regulating cargo transport to organize membrane domains at the cell surface.

**Key points:**

1. Arl15 primarily localizes to Golgi and plasma membrane, including filopodia
2. Membrane localization of Arl15 is dependent on Golgi integrity or Arf1 activation
3. Arl15 knockdown mislocalizes STX6-dependent Golgi localized cargo required for cell surface organization and reduces the filopodial number
4. Arl15 is involved in cell spreading, adhesion, and migration

## Introduction

Arf or Arf-like (Arl) GTPases belong to the Ras superfamily and play a critical role in physiological processes such as cell signalling, migration, membrane trafficking, and cytoskeleton reorganization (D’Souza-Schorey and Chavrier, 2006; Donaldson and Jackson, 2011; Sztul et al., 2019). Arl family includes 21 known Arl GTPases in human that regulate Golgi organization, vesicular transport, cytoskeletal assembly, and cilia formation (Burd et al., 2004; D’Souza-Schorey and Chavrier, 2006; Donaldson and Jackson, 2011; Sztul et al., 2019). Recent studies report the involvement of Arls in pathological conditions (Casalou et al., 2020). For example, Arl8B regulates prostate cancer in mice (Dykes et al., 2016); however at the cellular level, it controls membrane trafficking towards lysosomes by recruiting the HOPS complex (Khatter et al., 2015). Arl3, Arl6, and Arl13b play distinct roles in cilia biogenesis (Fisher et al., 2020; Li et al., 2012). The expression of Arl11 (ARLTS1) is altered in ovarian and breast cancers (Akisik et al., 2011; Petrocca et al., 2006), and regulates ERK signaling in macrophages (Arya et al., 2018). Arl4c has been implicated in colorectal cancer (Fujii et al., 2015) and modulates filopodium formation and cell migration through interactions with filamin-A (Chiang et al., 2017). Arl1 localizes to the *trans*-Golgi network (TGN) and regulates neuronal development, innate immunity, lipid droplet and salivary granule formation in addition to cargo transport (Yu and Lee, 2017). These studies indicate that Arls are involved in many cellular processes in addition to membrane trafficking. The function of Arl15 in regulating vesicular transport and the associated cellular processes are presently not well understood.

Genome-wide association studies delineated association of the genetic locus of *ARL15* gene is associated with multiple metabolic traits such as lower adiponectin and higher fasting insulin levels, increased risk of coronary heart disease and type 2 diabetes, insulin resistance and adiposity (Matsuba et al., 2016; Richards et al., 2009; Scott et al., 2014; Scott et al., 2012; Yaghootkar et al., 2014). Arl15 also regulates adipocyte differentiation and adiponectin secretion in murine 3T3-L1 cells and is mutated in lipodystrophy patients (Rocha et al., 2017). Depletion of Arl15 reduces insulin secretion in the human EndoC-βH1 cell line; these results indicate a strong link to diabetes (Thomsen et al., 2016). Additionally, Arl15 has been implicated in childhood obesity (Glessner et al., 2010), body shape (Ried et al., 2016), rheumatoid arthritis (Pandey et al., 2021; Sharma et al., 2020), and magnesium deficiency in insulin resistance and obesity (Corre et al., 2018). Zhao et al. showed that insulin stimulation enhanced the levels of Arl15 in C2C12 myotubes and possibly caused the activation of insulin signaling (Zhao et al., 2017). Shen and co-workers showed that high glucose-induced endothelial dysfunction in human umbilical vein endothelial cells (HUVECs) was attenuated by the enhanced expression of Arl15 (Shen et al., 2019). More recent studies show that Arl15 interacts with CNNM (cyclin M) proteins to inhibit the magnesium transport activity through enhancing their N-glycosylation (Zolotarov et al., 2021), and binding of Arl15 to Smad4 promote the Smad-complex formation and regulates TGFβ signaling (Shi et al., 2022). Overall, these studies place Arl15 as a signaling molecule in several genetic and metabolic traits in addition to magnesium homeostasis. The specific role of Arl15 in cargo trafficking is however not well understood.

Cell migration, invasion and adhesion are pivotal steps of many adherent cell types. Defects in any of these processes lead to various pathologies (Te Boekhorst et al., 2016). The role of Golgi localized Arf proteins, such as Arf1, Arf3 and Arf4, have been reported in migration, adhesion and cell proliferation (Howley et al., 2018; Huang et al., 2019; Luchsinger et al., 2018; Schlienger et al., 2015) in addition to their link to cancer progression (Casalou et al., 2016). The role of Arls, other than Arl4, in regulating cell adhesion or migration has however not been addressed.

In this study, we characterized the cellular function of Arl15. Extensive localization analysis in HeLa cells, using GFP-tagged Arl15, demonstrate that Arl15 predominantly localizes to the Golgi and plasma membrane (PM), including filopodia, and a cohort of tubular recycling endosomes. Treatment of stable cells expressing Arl15-GFP with small molecular inhibitors showed that the membrane localization of Arl15 was independent of the actin cytoskeleton but depended on the integrity of the Golgi. Further, Arf1 GTPase cycle possibly regulates the localization of Arl15 to Golgi but not the expression or its family GEFs or GAPs. Site-directed mutagenesis analysis identified a non-classical residue valine at the 80^th^ position in the GTP-binding domain of Arl15 which regulates its activity. Mutation at V80A mislocalize the Arl15 to cytosol and functions as a dominant-negative form which results in reduced number of filopodia. The siRNA-mediated depletion of Arl15 in HeLa cells reduced filopodial number, mislocalized Golgi localized cargo (caveolin-2 and STX6), altered the organization of focal adhesion sites, and in-and-out trafficking from the cell surface. At the cellular level, the depletion of Arl15 showed reduced cell migration and enhanced cell spreading. Traction force microscopy experiments demonstrate increased tractions in Arl15 depleted cells that had enhanced the number of focal adhesion puncta compared to control cells. Interestingly, biochemical adhesion assays showed that Arl15 activation was independent of integrin-mediated adhesion. These studies show that Arl15 regulates cargo trafficking from Golgi to organize the cell surface; this controls the formation of filopodia, cell migration, spreading and adhesion.

## Results

### Arl15 localizes primarily to Golgi and plasma membrane, including filopodia

Epitope-tagged Arl15 primarily localizes majorly to the Golgi and a cohort to the cell surface (Rocha et al., 2017; Shi et al., 2022; Wu et al., 2021; Zhao et al., 2017; Zolotarov et al., 2021). Here, we evaluated the reported Arl15 intracellular localization in various mammalian cell types. We tagged mouse Arl15 with GFP or mCherry at the C-terminus due to the unavailability of antibodies specific to localize endogenous Arl15. Note that both human and mouse Arl15 sequences are 97.1 % homologous to each other (**Supplementary Fig. 1A**). A comparison of human and mouse Arl15 sequences by homology modelling using AlphaFold2 displayed similar structural folds with a difference of six residues between the orthologs (**Supplementary Fig. 1A**). To reduce the variability in transgene expression, we prepared Arl15-GFP stably expressing HeLa cells using lentivirus-mediated transduction (referred to here as HeLa:Arl15-GFP stable cells). Fluorescence microscopy analysis of stable cells displayed uniform expression of Arl15-GFP (data not shown). We used a non-clonal population of cells throughout this study. Immunoblotting analysis confirmed the expression of Arl15-GFP with a partial GFP cleavage in the HeLa:Arl15-GFP stable cells (**Supplementary Fig. 1B**).

Immunofluorescence microscopy (IFM) of Arl15-GFP expressing HeLa cells with organelle-specific markers revealed that Arl15-GFP predominantly localizes to the PM and extracellular protrusions such as filopodia (**Fig. 1**). Consistently, Arl15-GFP showed colocalization with a PM marker, wheat germ agglutinin (WGA)-Alexa Fluor 594 (**Fig. 1A**). Analysis of HeLa:Arl15-GFP stable cells with phalloidin Alexa Fluor 594 (stains F-actin), or expressed with mCherry-UtrCH to mark actin, showed significant colocalization of Arl15-GFP with labeled phalloidin or mCherry-UtrCH; these results suggest that Arl15 mainly localizes to PM and filopodial protrusions (**Fig. 1B** and **1C**). Intracellularly, Arl15-GFP localized to Golgi stacks near the perinuclear area and to long tubular structures resembling recycling endosomes (REs). Consistently, IFM analysis revealed that Arl15-GFP colocalized with *cis*- and *trans*-Golgi proteins GM130 and Golgin p230, respectively, and recycling endosomal marker KIF13-YFP (Shakya et al., 2018) (**Fig. 1D, 1E** and **Supplementary Fig. 1C**). Arl15-GFP did not exhibit localization to the endoplasmic reticulum (ER), early endosomes, lysosomes or focal adhesion sites marked by calnexin, EEA1, LAMP-1 and vinculin, respectively, during the colocalization experiments in HeLa cells expressing Arl15-GFP (**Supplementary Fig. 1C**). We also did not observe any significant difference in cell size, as measured by FACS, between HeLa and HeLa:Arl15-GFP stables (**Supplementary Fig. 1D**). To test whether the localization of Arl15 was conserved in other mammalian cell types, we expressed Arl15-GFP transiently in A549, primary human keratinocytes and Neuro 2a cells. These results show that similar to HeLa cells, Arl15-GFP localizes to PM along with filopodia and Golgi in all mammalian cell types (**Fig. 1F**). These data hence confirm the localization of Arl15-GFP primarily to Golgi, PM (including filopodia), and a cohort to REs and is conserved in different mammalian cell types.

**Figure 1.**
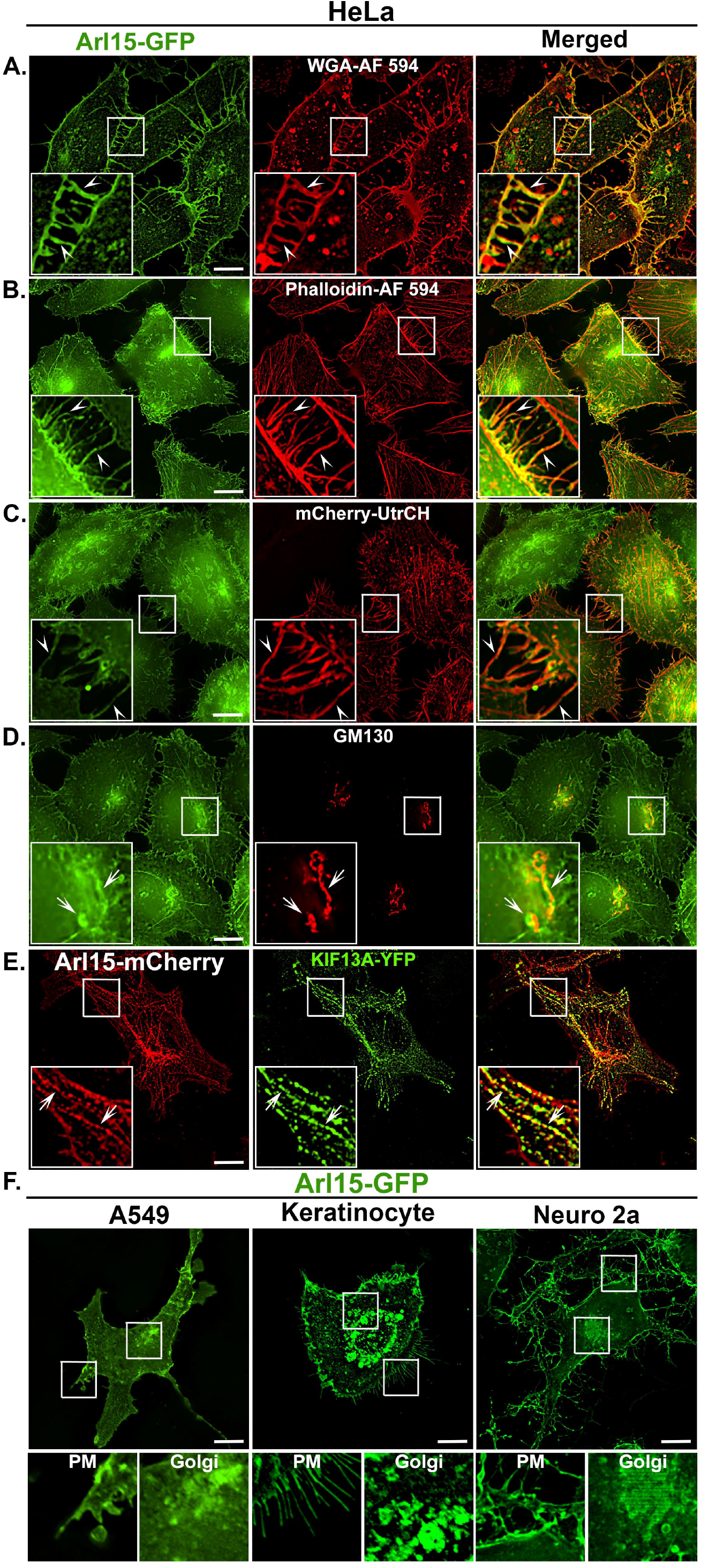
Arl15-GFP localizes to the plasma membrane, filopodia, recycling endosomes and Golgi. IFM analysis of HeLa:Arl15-GFP stable cells. Cells were stained with WGA-Alexa Fluor 594 (A), phalloidin-Alexa Fluor 594 (B), transfected with mCherry-UtrCH (C) or stained with anti-GM130 antibody (D) and then imaged. In (E), plain HeLa cells were co-transfected with Arl15-mCherry and KIF13-YFP. Arrowheads point to the localization of Arl15-GFP to PM, including filopodia. Arrows point to Golgi or KIF13A-positive recycling endosomes. (F) IFM analysis of Arl15-GFP expressing A549, primary human keratinocytes and Neuro 2a cells. Insets are magnified views of the white boxed areas. In F, the magnified view of insets for both PM and Golgi are shown separately. Scale bars, 10 µm.

### Cytoskeleton assembly does not modulate the localization of Arl15 to the plasma membrane

Studies have shown that Arl4A (Patel et al., 2011) and Arl4D (Li et al., 2007) regulate actin cytoskeleton remodelling; Arl13B interacts with actin (Barral et al., 2012), and Arl2 participates in tubulin assembly for microtubule network (Francis et al., 2017). To test the possible recruitment of Arl15 to the PM (also to filopodia) and its dependence on cytoskeleton network/assembly, we transfected stable HeLa:Arl15-GFP cells with mCherry-UtrCH (labels actin cytoskeleton) and treated with latrunculin A to inhibit actin polymerization. Cells appear to shrink due to actin depolymerization after 15 min of treatment; the localization of Arl15-GFP to PM however remained unchanged as compared to DMSO treated cells (**Fig. 2A**). Cells displayed normal size with Arl15-GFP on PM along with filopodia upon rescue of the cells by removing latrunculin A, and following incubation in a complete medium for 4 h to restore the actin assembly (**Fig. 2A**). Because Rho GTPases, such as Cdc42 and Rac1, play a key role in local F-actin organization and support the formation of filopodia and lamellopodia, we tested if the membrane recruitment of Arl15 was regulated by actin remodeling (Krugmann et al., 2001; Steffen et al., 2013). We treated mCherry-UtrCH expressing HeLa:Arl15-GFP stable cells with a small molecular inhibitor of Rac1 (CAS 1177865-17-6) or Cdc42 (ML141) for 1 h. Interestingly, these inhibitors showed no effect on Arl15-GFP localization to PM/Golgi (**Fig. 2B**) but displayed fewer filopodia (data not shown). We next tested the role of integrin-dependent focal adhesion kinase (FAK), an upstream factor of Cdc42/RhoA/Rac1 GTPases (Nobes and Hall, 1995), in the membrane recruitment of Arl15 using FAK inhibitor (PF-573228) for 1 h. Compared to control cells, FAK inhibitor-treated cells were more spread out and flattened with prominent extracellular protrusions without affecting Arl15-GFP localization to PM/Golgi (**Fig. 2C**). Overall, these studies indicate that neither actin depolymerization nor the key regulatory molecules of filopodia/lamellipodia affect the localization of Arl15 to PM/Golgi. We next tested the role of Golgi stacking/assembly in maintaining the localization of Arl15 to Golgi/PM. We treated the HeLa:Arl15-GFP stables with nocodazole for 90 min to disrupts microtubule assembly and stained cells for *cis*-Golgi protein GM130. Nocodazole treatment disassembled the Golgi stacks into small vesicles (Presley et al., 1998), which mislocalized Golgi pool of Arl15 without affecting its PM localization (**Fig. 2D**). These findings suggest that the membrane association of Arl15 is independent of the actin cytoskeleton but may depend on the Golgi organization.

**Figure 2.**
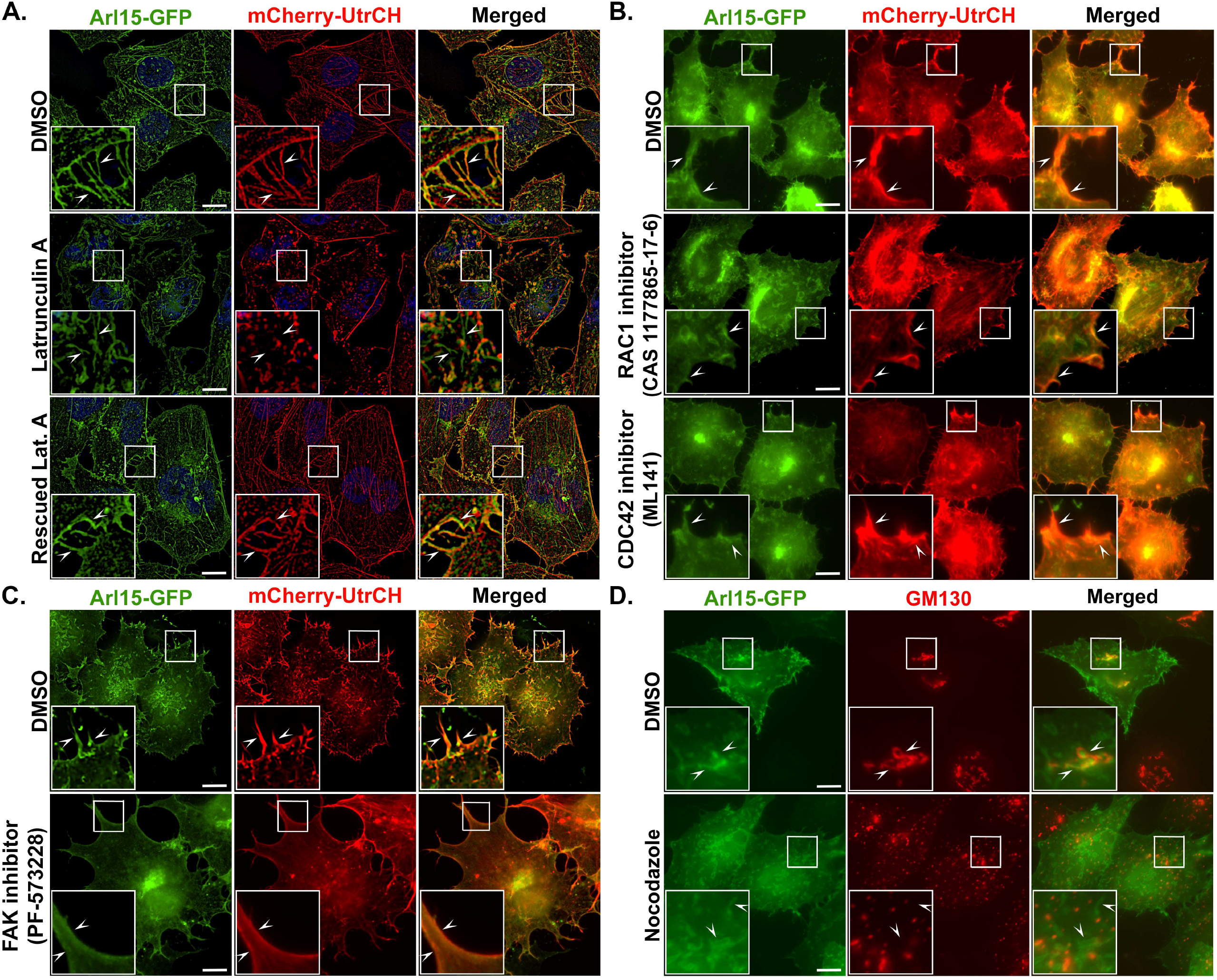
Cytoskeleton assembly do not regulate the localization of Arl15-GFP to PM. IFM analysis of HeLa:Arl15-GFP stable cells treated with inhibitors of actin polymerization (latrunculin A for 15 min, in A), RAC1 (CAS 1177865-17-6 for 1 h, in B), CDC42 (ML141 for 1 h, in B), FAK1 (PF-573228 for 1 h, in C), or microtubule assembly/disassembly (nocodazole for 1.5 h, in D) separately. Cells were treated with DMSO as a control in all experiments. In A – C, cells were transfected with mCherry-UtrCH to label the actin cytoskeleton. In D, cells were stained with anti-GM130 antibody. In A, cells were washed post inhibitor treatment and then rescued for 4 h in complete medium (rescued latrunculin A). Arrowheads point to the localization of Arl15-GFP to PM, including filopodia (A-C). In D, arrowheads represent Golgi and/or PM localization of Arl15-GFP. Insets are magnified views of the white boxed areas. Scale bars, 10 µm.

### Golgi integrity/stacking controls the membrane association of Arl15

Arl15 localizes to Golgi stacks as reported for other Arfs/Arls (Arf1, 3, 4, 5 and Arl1, 3, 5, 4A) (Marwaha et al., 2019; Sztul et al., 2019). To test whether Golgi integrity or stack assembly is required for the membrane association of Arl15, we treated the HeLa:Arl15-GFP stables or plain HeLa cells expressing Arf1-GFP (positive control) or GFP (negative control) with brefeldin A (1.25 µg/ml) for 24 h and then fixed and stained for Golgi protein, GM130. Note that the positive control Arf1-GFP is known to regulate the Golgi assembly/stacking/integrity (Donaldson et al., 2005). GM130 stained Golgi in GFP-expressing HeLa cells showed puncta and were dispersed throughout the cytosol following brefeldin A treatment as compared to DMSO (**Fig. 3A**). Arf1-GFP showed cytosolic localization after brefeldin A treatment of HeLa cells. Surprisingly, Arl15-GFP localized to the cytosol as similar to Arf1-GFP in GM130 dispersed cells that are indicative of brefeldin A treatment in HeLa cells (**Fig. 3A**). We used another small molecular inhibitor of GBF1 (nucleotide exchange factor of Arf1), golgicide A, known to alter the Golgi integrity, to confirm these results (Sáenz et al., 2009). These studies showed that treatment of the cells with golgicide A (20 µg/ml) for 24 h resulted in mislocalization of Arl15-GFP to the cytosol and dispersed the GM130-positive Golgi puncta (**Fig. 3A**). Arf1-GFP also localized to cytosol upon treatment of HeLa cells with golgicide A (**Fig. 3A**). Overall, these studies indicate that the localization of Arl15 to Golgi/PM is dependent on Golgi integrity/stacking mediated through Arf1 activity.

**Figure 3.**
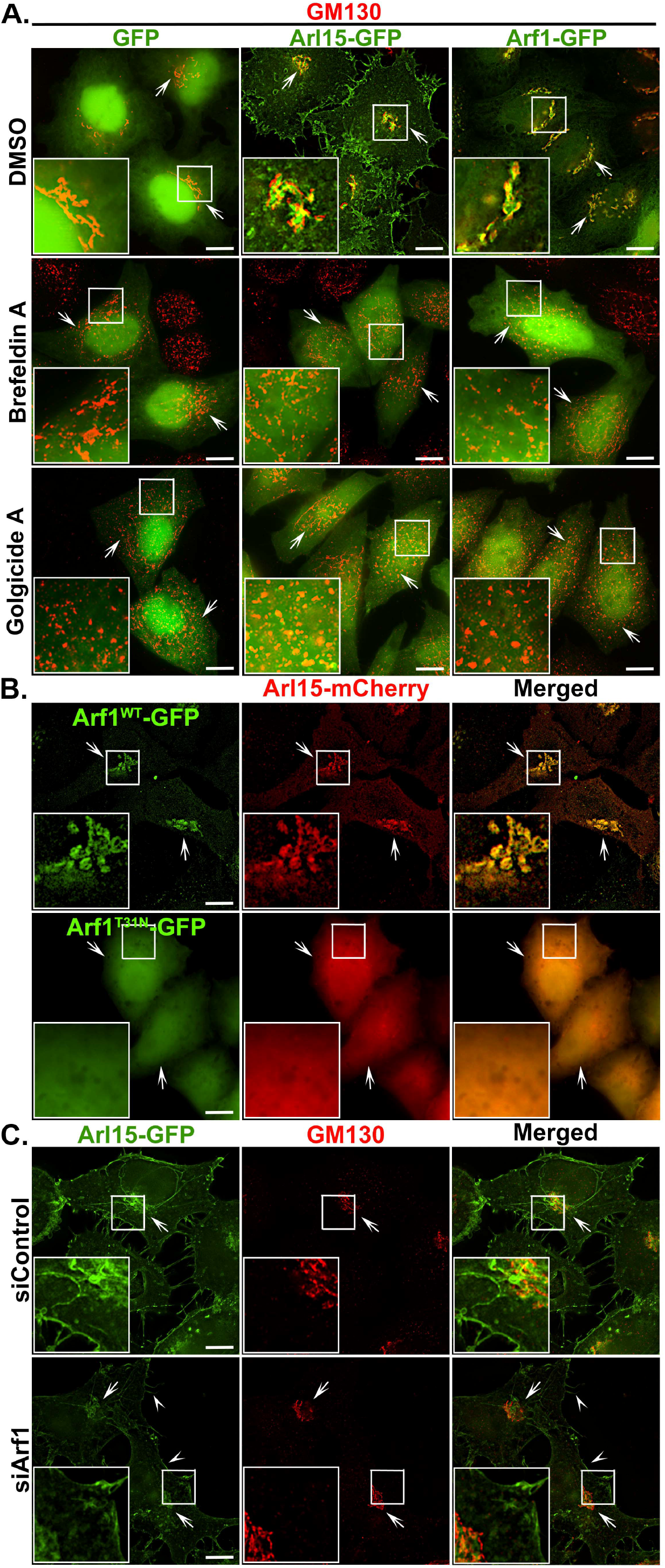
Golgi integrity through Arf1 activity regulates the membrane localization of Arl15-GFP. (A) IFM analysis of HeLa cells expressing GFP, Arl15-GFP or Arf1-GFP. Cells were treated with DMSO (as control), brefeldin A or golgicide A for 24 h followed fixation, staining with anti-GM130 antibody and then imaged. Note: Treating cells with brefeldin A or golgicide A mislocalizes both Arf1 and Arl15 to the cytosol. Arrows point to the intact Golgi in DMSO treated cells and dispersed Golgi in drug-treated cells. (B) IFM analysis of HeLa cells expressing Arl15-mCherry with Arf1^WT^-GFP or Arf1^T31N^-GFP. Arrows point to Golgi or cytosolic localization of the proteins. (C) IFM analysis of siControl and siArf1 treated HeLa:Arl15-GFP stable cells. Cells were stained with anti-GM130 antibody. Arrows point to Golgi, and arrowheads point to the PM (including filopodia) localized Arl15-GFP. Insets are magnified views of the white boxed areas. Scale bars, 10 µm.

### GTPase cycle of Arf1 maintains the localization of Arl15 to Golgi/PM

Arf1 localizes to entire Golgi stacks (*cis, medial* and *trans*), as well as to the ER-Golgi intermediate compartments (ERGIC), recycling endosomes (REs) and a cohort to PM (Donaldson and Jackson, 2011). The GTPase activity of Arf1 also plays a vital role in maintaining Golgi integrity and function (Donaldson et al., 2005; Donaldson and Klausner, 1994; Jackson, 2018; Lippincott-Schwartz et al., 1998; Yu and Lee, 2017). Because small molecules, such as brefeldin A/golgicide A, affect the Arf1 activity and mislocalize the Arl15 to the cytosol (Presley et al., 1998; Sáenz et al., 2009), we tested if Golgi localized Arf1 directly regulates the membrane localization of Arl15. Coexpression of Arf1^WT^-GFP and Arl15^WT^-mCherry in HeLa cells showed a significant overlap in colocalization between these small GTPases at Golgi (**Fig. 3B**). The expression of a dominant-negative form of Arf1^T31N^-GFP with Arl15^WT^-mCherry results in mislocalization of Arl15 to the cytosol (**Fig. 3B**); these results suggest that active Arf1 is required for Arl15 localization to the membrane.

We evaluated the importance of Arf1 expression in Arl15 localization. SiRNA-mediated knockdown of Arf1 in HeLa cells showed reduced levels of endogenous Arl15 (**Supplementary Fig. 2A**). In contrast, Arl15 depletion using gene-specific siRNA did not alter endogenous Arf1 levels (**Supplementary Fig. 2A and 2B**); these results suggest a role for Arf1 in maintaining Arl15 expression. Arf1 knockdown in stably expressing Arl15-GFP HeLa cells did not display any change in the Arl15 localization from Golgi/PM (**Fig. 3C**). Arf1 depletion did not also affect Golgi integrity/stacking in HeLa cells (**Fig. 3C**). These data suggest the presence of compensatory mechanisms to maintain the Golgi function may be mediated by Arf3, 4 and 5 in HeLa cells (Pennauer et al., 2022). To explore the role of Arf1 function up/downstream of Arl15, we studied the localization of Arf1-GFP in Arl15 knockdown HeLa cells. IFM studies showed that Arl15 knockdown did not affect Arf1 localization to Golgi or its expression in the cells (**Supplementary Fig. 2A and 2C**). These results demonstrate that the Arf1 activity alone (active/inactive), but not the expression level, may play a significant role in the membrane association of Arl15; Arf1 likely acts upstream of Arl15.

### Non-classical GTPase domain Arl15^V80A^ mutant acts as a dominant-negative form and reduces the filopodial number

To test the role of GTPase cycle and/or myristoylation of Arl15 in regulating its localization to Golgi/PM, we mutated the conserved residues in Arl15 using site-directed mutagenesis (**Fig. 4A**). Alignment of Arl15 amino acid (aa) sequence with other Arf and Arl family GTPases revealed that Arl15 showed the closest homology with Arl2 (UniProt: P36404) and Arf6 (UniProt: P62330) with approximately 34% sequence identity (data not shown). The alignment of Arl15 with the crystal structure of Arl2 [PDB: 3DOE] generated a homology structure of Arl15 having a GTP binding pocket. The obtained structure displayed the residues G39, L40, T41, G42, S43, G44, K45, T46, E82, L83 G85, N142, H143, Q144, D145, and K146 surrounding the GTP binding domain of Arl15 (**Fig. 4B**). To identify the key conserved residues that act as classical constitutive active (Q) and dominant residues (S/T) in Arl15 (Kahn et al., 2006; Pasqualato et al., 2002), we analysed the Arl15 GTP-binding domain. The analysis identified non-classical residue alanine at 86^th^ position (A86) instead of Q (glutamine), which is conserved in other Arl/Arf GTPases. We changed the residue A86 into classical Q86 or constitutive active L86 form in the Arl15 predicted structure; these mutations did not however alter the GTP-binding domain grossly (**Fig. 4B**).

**Figure 4.**
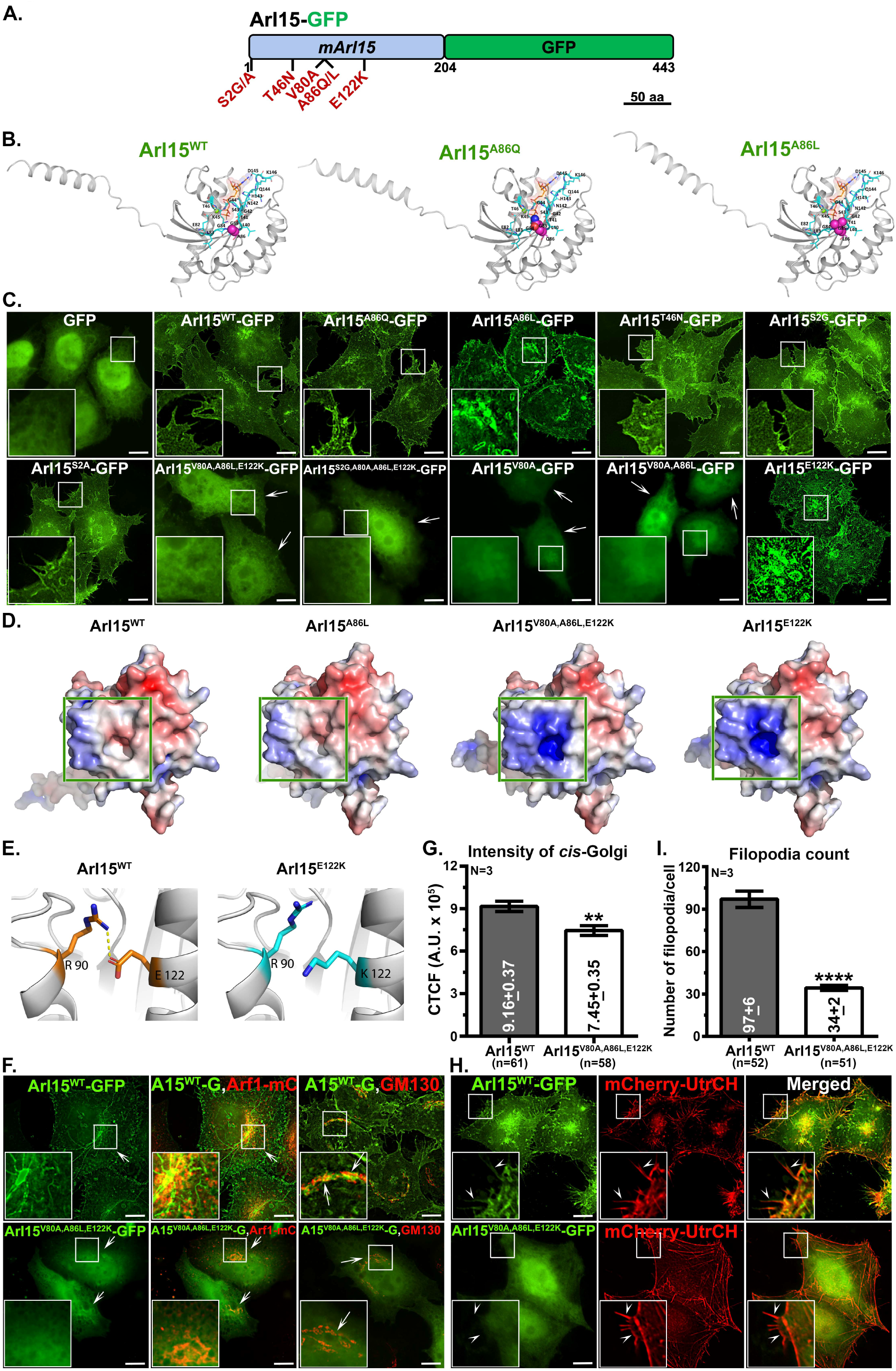
Mutational analysis identified non-classical mutation V80A in the GTP-binding domain of Arl15 as a dominant-negative form. (A) A schematic representation of mouse Arl15-GFP has been drawn with an indicated scale bar. Mutations at the myristoylation site (S2G or S2A), GTPase domain (classical mutations T46N or A86L and non-classical residues V80A, A86Q) and on the surface (E122K) are generated and used in the study. (B) The predicted structural view of Arl15^WT^, Arl15^A86Q^ and Arl15^A86L^ was generated through homology modelling. The residues involved in GTP binding are highlighted in cyan and the GTP is highlighted in orange. (C) IFM analysis of HeLa cells expressing GFP, Arl15^WT^-GFP or different mutants of Arl15-GFP as indicated in the figure. Arrows point to cytosolic localization of Arl15-GFP. (D) Predicted surface electrostatics comparison of Arl15^WT^ with its mutants Arl15^A86L^, Arl15^V80A,A86L,E122K^ and Arl15^E122K^ generated using APBS plotted in the range from −10 to +10 to −5 to +5 *k*_*B*_*Te*^*-1*^. Green boxes represent the change in surface charge in mutants Arl15^V80A,A86L,E122K^ and Arl15^E122K^ compared to Arl15^WT^ or Arl15^A86L^. (E) Schematic representation of a salt bridge between R90 and E122 residues in Arl15^WT^ and its disruption in Arl15^E122K^ mutant. (F, H) IFM analysis of HeLa cells expressing Arl15^WT^-GFP or Arl15^V80A,A86L,E122K^-GFP. Cells are either co-transfected with Arf1-mCherry (F) or mCherry-UtrCH (H) or stained with anti-GM130 antibody (F). Arrows point to the localization of Arl15^WT^-GFP to Golgi or cytosol (F). Arrowheads point to the localization of Arl15^WT^-GFP to PM, including filopodia (H). In all images, insets are magnified views of the white boxed areas. Scale bars, 10 µm. (G, I) Plots represent the intensity (CTCF) of *cis*-Golgi (G) and filopodia count/cell (I) in HeLa cells expressing Arl15^WT^-GFP or Arl15^V80A,A86L,E122K^-GFP. The average values in mean ± s.e.m. are indicated on the graph. N, the number of repeats and n, the total number of cells. \*\**p*≤0.01and \*\*\*\**p*≤0.0001.

To understand the functionality of these mutations, we generated Arl15^A86Q^-GFP and Arl15^A86L^-GFP mutants and expressed them in HeLa cells. These mutants localized to Golgi/PM as similar to Arl15^WT^-GFP, but did not alter the Golgi structure (**Fig. 4C**). We next mutated T46 to N46 (threonine to asparagine) in Arl15, a classical dominant-negative mutation observed in the majority of Arfs/Arls. The Arl15^T46N^-GFP mutant did not display the expected cytosolic localization but localized as similar to the wild type (**Fig. 4C**). Thus, Arl15^T46N^ does not act as a dominant-negative form like other Arls. The second position in many Arls is glycine (Kahn et al., 2006; Pasqualato et al., 2002), which is generally myristoylated and facilitates the anchoring of Arls to the membranes (Gillingham and Munro, 2007). Interestingly, Arl8 and Arf-related protein 1 (ARFRP1, yeast homologue Arl3p) do not contain glycine residue, and these GTPases are shown to anchor to the membranes through acetylation (Behnia et al., 2004; Hofmann and Munro, 2006; Setty et al., 2004). In contrast, Arl2 possess myristoylation motif but was reported not to get myristoylated (Sharer et al., 2002). Analysis of the myristoylation motif revealed that Arl15 contains serine (S) at +2 in place of glycine (G), which is found in other Arf/Arl GTPases. We tested if the conversion of S to G (S2G) altered the localization of Arl15-GFP in HeLa cells. The expression of Arl15^S2G^-GFP in HeLa cells showed its localization to Golgi/PM which was similar to wild type Arl15 (**Fig. 4C**). We next prepared Arl15^S2A^-GFP to evaluate the role of serine at 2^nd^ position, which showed similar localization as Arl15^WT^-GFP in HeLa cells (**Fig. 4C**). The additive mutation of S2G with A86L in Arl15-GFP also showed no effect on its localization (data not shown). These analyses suggest that S2 residue may not play a key role in the membrane recruitment of Arl15. During the screening of different plasmids having Arl15^A86L^ mutation, we identified a form showing cytosolic localization in HeLa cells. The DNA sequence analysis of the construct showed two additional mutations at V80A and E122K in addition to A86L in Arl15. An additive mutation of S2G to this Arl15^V80A,A86L,E122K^ mutant showed cytosolic localization; these data indicate that the myristoylation-positive motif does not compensate the triple mutation. Analysis of V80A, A86L and E122K mutations in the predicted Arl15 structure showed a change in surface hydrophobicity in the triple mutant compared to Arl15^A86L^ or Arl15^WT^ (**Fig. 4D**). Homology modelling analysis further revealed that the surface hydrophobicity is contributed majorly by E122K mutation in Arl15 (**Fig. 4D**), due to a loss of salt bridge formation (between E122 and R90 residues) on the surface (**Fig. 4E**). We also hypothesized that the mutation at V80A located in the GTP-binding domain of Arl15 possibly alters the GTPase activity. To test if the loss of salt bridge (E122K) or GTPase activity (V80A) contributes to the cytosolic localization of Arl15, we studied the localization of these mutants in HeLa cells. Individual mutations in Arl15 displayed Arl15^V80A^ as cytosolic localization compared to Arl15^A86L^ or Arl15^E122K^ mutants, localized as similar to Arl15^WT^ (**Fig. 4C**). Note that few cells expressing Arl15^V80A^-GFP showed small clusters (presumably Golgi) in the perinuclear region along with cytosolic localization (**Fig. 4C**). A double mutant containing V80A and A86L (predicted constitutive active form) did not rescue the cytosolic localization of Arl15^V80A^ (**Fig. 4C**). These data suggest a novel role of V80A in converting the Arl15 into dominant-negative form, possibly due to a defect in the binding of GTP. However, we continued with Arl15^V80A,A86L,E122K^ cytosolic mutant to study the functionality of Arl15 in HeLa cells. Overall, site-directed mutagenesis analysis in Arl15 identified a unique mutation at V80A in the GTP-binding domain, which regulates its localization to Golgi/PM.

Based on these results, we hypothesized that the cytosolic Arl15^V80A,A86L,E122K^ mutant may act as a dominant-negative form and affects the cellular function. We expressed Arl15^V80A,A86L,E122K^-GFP (cytosolic mutant) in the HeLa cells and studied its effects on Golgi integrity and the status of filopodia in comparison with Arl15^WT^-GFP. The cytosolic mutant of Arl15 showed no effect on the localization of Arf1-mCherry or GM130 to Golgi compared to Arl15^WT^ (**Fig. 4F**). Quantification of GM130 intensity (indicative of *cis*-Golgi stacks measured by corrected total cell fluorescence, CTCF) in the cells showed a reduced number (0.81 folds) with mild dispersion in the cells expressing Arl15^V80A,A86L,E122K^-GFP mutant compared to Arl15^WT^-GFP in HeLa cells (**Fig. 4F and 4G**). These results indicate that Arl15 may not regulate the Golgi organization or localization of Arf1 to Golgi. The visualization of filopodia by IFM using mCherry-UtrCH displays a dramatic difference in filopodia pattern, number, and organization in cells expressing Arl15^V80A,A86L,E122K^-GFP mutant compared to Arl15^WT^-GFP (**Fig. 4H**). We also observed mildly enhanced stress fibers in cells expressing cytosolic mutant as compared to wild type Arl15 (**Fig. 4H**). A visual quantification of filopodial number per cell showed a significant reduction (to 0.35 folds) in the cells expressing Arl15^V80A,A86L,E122K^-GFP mutant compared to Arl15^WT^-GFP (**Fig. 4I**). These studies in a nutshell demonstrate that Arl15^V80A,A86L,E122K^ mutant acts as dominant-negative form and alters the filopodial biogenesis (length and number) in HeLa cells.

### Arl15 depletion mislocalizes Golgi localized cargoes and alters cell surface transport and organization

We performed a siRNA-mediated knockdown of Arl15 in HeLa cells to understand its molecular function. Immunoblotting analysis showed a dramatic decrease in Arl15 levels in cells transfected with gene-specific siRNA (siArl15 or siA15) as compared to control siRNA (siControl or siCont.) (**Supplementary Figure 2B**). Note that the currently available commercial antibody (11934-1-AP, Proteintech) works only for western blotting but is unsuitable for IFM studies. We carried out the knockdown of Arl15 using 3’-UTR-specific siRNA (siArl15^3-UTR^) in HeLa:Arl15-GFP stables (**Supplementary Figure 2B**) to test the antibody specificity. Immunoblotting results showed reduced levels of endogenous Arl15 detected with the commercial antibody without affecting the Arl15-GFP transgene expression (**Supplementary Figure 2B**). We next analysed the status of Golgi in Arl15 depleted HeLa cells by studying the localization of structural and cargo proteins. IFM analysis of Golgi localized tethering proteins such as GM130 and GRAPS65 showed no observable change in their localization other than partial dispersal of Golgi in Arl15 knockdown as compared to control cells (**Fig. 5A**). Quantification of GM130 fluorescence intensity (CTCF) showed a slight increase in siArl15 compared to siControl cells (**Fig. 5B**). Arl15 depletion in HeLa cells did not affect Arf1 expression or Arf1-GFP localization to Golgi other than its dispersion (**Supplementary Fig. 2A and 2C**). Interestingly, Arl15 knockdown in HeLa cells displayed mislocalization of caveolin-2 (Cav-2) from Golgi (**Fig. 5A**) (Mora et al., 1999). The number of cells showing Golgi localized caveolin-2 staining correspondingly decreased in addition to reduced protein expression in siArl15 compared to siControl cells (**Fig. 5C and 5D**). In contrast, Arl15 knockdown cells showed no effect on the localization of caveolin-1 compared to control cells (**Supplementary Fig. 2D**). Arl15 depleted cells also showed reduced levels of Golgi localized SNARE syntaxin 6 (STX6) compared to control cells by IFM and immunoblotting studies (**Fig. 5A, 5B and 5D**) (Bock et al., 1997). To test whether the mislocalized Cav-2 and STX6 are targeted to lysosomes in siArl15 cells, we treated the cells with bafilomycin A1 (25 nM for 6 h) along with control cells. Surprisingly, Cav-2 and STX6 rescued their Golgi localization without accumulating in lysosomes after the treatment of siArl15 cells with bafilomycin A1 (**Supplementary Fig. 2E**). Note that Arl15 knockdown in HeLa cells showed no effect on LAMP-1-positive lysosomes compared to control cells (**Supplementary Fig. 2D**). Immunoblotting analysis showed Cav-2 and STX6 protein levels were not restored in siArl15 cells equivalent to siControl cells after the treatment with bafilomycin A1 or MG132 (**Supplementary Fig. 2F**). These results suggest different possibilities such as (1) the 6 hr treatment with bafilomycin A1 was not sufficient to enrich these proteins in the lysosomes; (2) bafilomycin A1 treatment may have blocked the transport of these molecules to lysosomes; or (3) the newly biosynthesized molecules possibly accumulated in the Golgi of siArl15 cells, which requires future investigation.

**Figure 5.**
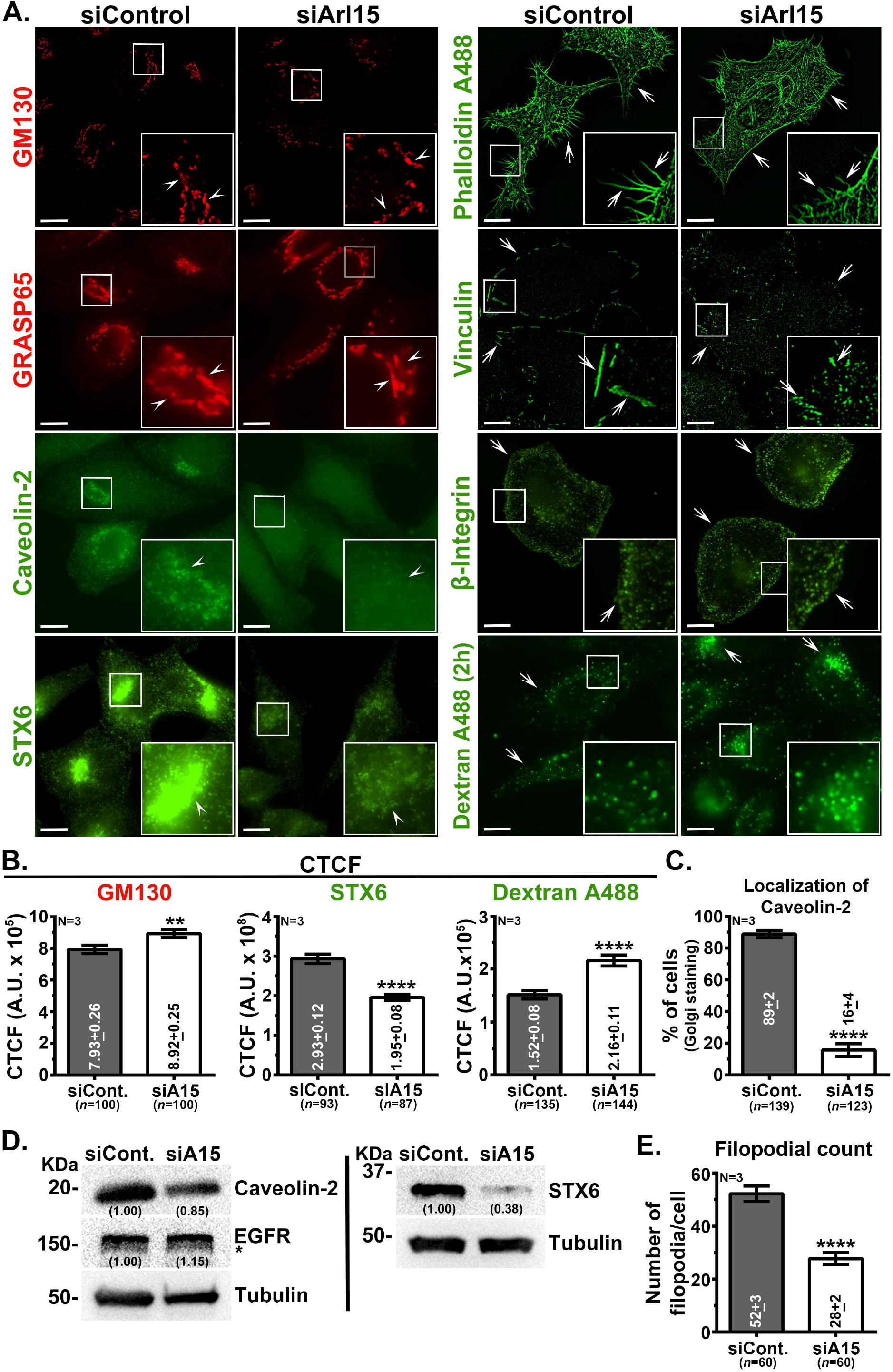
Arl15 depletion in HeLa cells mislocalizes cargo of Golgi, reduces filopodia count and alters cell surface trafficking. (A) IFM analysis of siControl and siArl15 treated HeLa cells. In the left panel, cells were stained for Golgi-associated proteins/cargo. Arrowheads point to the localization of proteins to Golgi. In the right panel, cells were stained for the actin cytoskeleton, cell surface proteins or internalized with soluble cargo. Arrows point to the filopodia, cell surface proteins or internalized cargo. In all images, insets are magnified views of the white boxed areas. Scale bars, 10 µm. (B) Plots represent the intensity (CTCF) of GM130, STX6 or internalized dextran Alexa Fluor 488 in siControl and siArl15 treated HeLa cells, arbitrary units (A.U.). The average values in mean ± s.e.m. are indicated on the graph. (C, E) Plots represent the percentage of Golgi localized caveolin-2-positive cells (C) or the number of filopodia/cell (E) in siControl and siArl15 treated HeLa cells. The average values in mean ± s.e.m. are indicated on the graph. N, the number of repeats and n, the total number of cells. \*\**p*≤0.01 and \*\*\*\**p*≤0.0001. (D) Immunoblotting analysis of siControl and siArl15 treated HeLa cells for the expression of caveolin-2, EGFR and STX6. γ-tubulin is used as a loading control. *, the non-specific band developed with the antibody. The normalized fold change in the expression with respect to control is indicated on the blots.

We next tested the effect of Arl15 knockdown on PM organization/transport and filopodial biogenesis. Staining of the actin cytoskeleton with phalloidin Alexa Fluor 488 showed a decreased number of filopodia on the cell surface of Arl15 depleted HeLa cells compared to control cells (**Fig. 5A**). The visual quantification of filopodia displayed a significant decrease in the number of filopodia/cell (to 0.54 fold) in siArl15 compared to siControl cells (**Fig. 5E**). The localization of focal adhesion site markers such as vinculin and β1-integrin showed a defect in vinculin organization (appeared as punctate structures) on the cell surface without affecting the localization of β1-integrin in siArl15 cells compared to siControl cells (**Fig. 5A**). These results suggest a role for Arl15 in the assembly of focal adhesion sites (see below). We evaluated if the depletion of Arl15 alters the cell surface dynamics, such as endocytosis/recycling of cargo. The localization and expression of clathrin-mediated endocytosis cargo EGFR (traffics from the cell surface to lysosomes) showed no difference in siArl15 and siControl cells (**Supplementary Fig. 2D and Fig. 5D**). However, siArl15 cells showed enhanced uptake of pinocytosis marker dextran Alexa Fluor 488 and receptor-mediated endocytosis marker transferrin Alexa Fluor 594 (at 20 min) compared to control cells (**Fig. 5A, 5B, Supplementary Fig. 2G** and **2H**). The recycling rate of transferrin receptor (from endosomes, measured using Tf Alexa Flour 594 at 40 min chase) to the cell surface was moderately reduced (0.79 folds) in siArl15 compared to siControl cells (**Supplementary Fig. 2G** and **2H**). The cell surface lectin levels (measured using WGA Alexa Flour 594) were also reduced (0.86 folds) in siArl15 compared to siControl cells (**Fig. 2I**). These studies overall highlight a possible role for Arl15 in regulating cargo trafficking from the Golgi to the PM for the organization of cell surface membrane domains; those may regulate the filopodial number and endocytic/recycling cargo dynamics.

### Arf1 or Arf family-GEFs/GAPs do not modulate the activity of Arl15

We predict that Arl15 but not Arf1 or Arf family guanine exchanges factors (GEFs)/GTPase activating proteins (GAPs) modulate the trafficking of caveolin-2/STX6 from Golgi. We tested this hypothesis by performing the knockdown of individual genes Arf1; GBF1, BIG1, BIG2 (Arf family GEFs); ASAP1 or ASAP2 (Arf family GAPs) in HeLa cells (Casalou et al., 2020; Donaldson and Jackson, 2011; Gillingham and Munro, 2007; Sztul et al., 2019) and then studied the localization of caveolin-2/STX6 by IFM. The BIG2 gene depletion in HeLa cells was validated using IFM (data not shown) and the other genes by immunoblotting (**Supplementary Fig. 3A**). We initially analyzed the cells for the localization of Arl15-GFP to Golgi/PM and the cells displayed no change in membrane localization (**Fig. 6A** and **6B**). The pooled GEFs or GAPs knockdown in HeLa cells also did not mislocalize the Arl15-GFP to the cytosol (**Supplementary Fig. 3B**). The expression level of Arl15 was not reduced in any of the Arf family-GEFs/GAPs depleted cells except in Arf1 knockdown cells (**Fig. 6C and 6D**). These results suggest that the Arf family-GEFs/GAPs may not regulate the activity of Arl15. We further tested the localization of Golgi-associated proteins such as GM130 and GBF1, and Golgi cargo caveolin-2 and STX6 in the Arf1- or Arf family-GEFs/GAPs knockdown cells. The knockdown cells displayed no gross change in the localization of GM130 and GBF1 to Golgi (except GBF1 staining in siGBF1) (**Fig. 6A, 6B** and **Supplementary Fig. 3C**). Similarly, the localization and expression of caveolin-2 or STX6 to Golgi was grossly unaffected in the Arf1- or Arf family-GEFs/GAPs knockdown cells (**Fig. 6A – 6D**). We also observed reduced staining of caveolin-2 in siBIG2 cells and lower expression of STX6 in siArf1 or siBIG1 cells (**Fig. 6A – 6D**). A recent study described an interaction between ASAP2 and Arl15 in C2C12 myocytes and predicted that ASAP2 possibly acts as a GAP for Arl15 (Zhao et al., 2017). However, we have not observed any significant change in Arl15 and its dependent cargo (caveolin-2 and STX6) localization to Golgi in siASAP2 cells (**Fig. 6B and Supplementary Fig. 3D**). These results indicate that Arl15 recruitment to the Golgi is independent of Arf family-GEFs/GAPs expression but possibly dependent on the Arf1 GTPase cycle.

**Figure 6.**
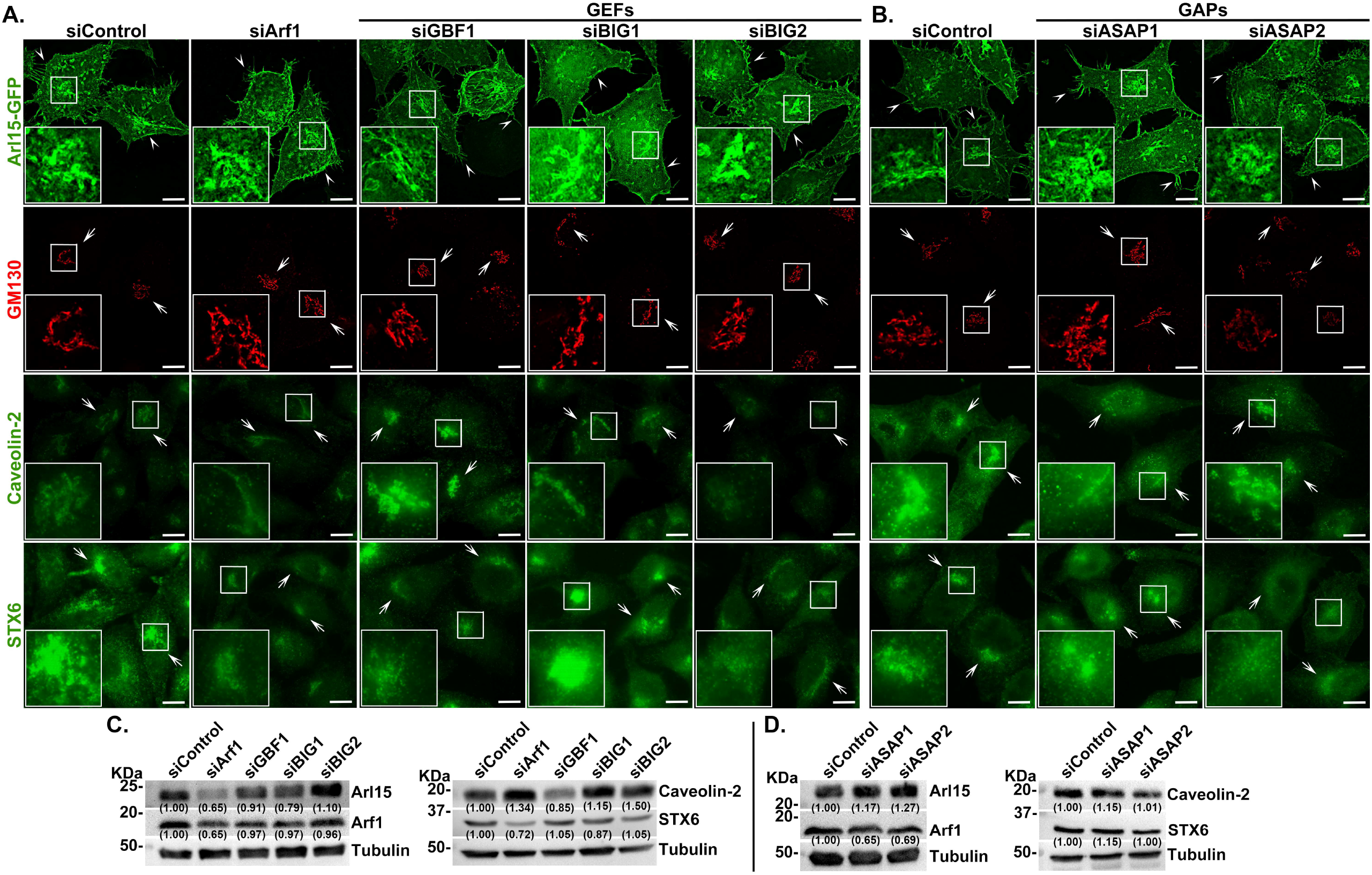
Arf family GEFs or GAPs do not regulate the membrane localization of Arl15-GFP or its cargo. (A, B) IFM analysis of siControl, siArf1, siArf family-GEFs (GBF1, BIG1, BIG2) and siArf family-GAPs (ASAP1, ASAP2) treated HeLa cells. Cells were stained for GM130, caveolin-2 or STX6. Arrowheads point to the localization of Arl15-GFP to PM, including filopodia. Arrows point to the localization of proteins to Golgi. In all images, insets are magnified views of the white boxed areas. Scale bars, 10 µm. (C, D) Immunoblotting analysis of HeLa cells treated with different siRNAs separately as indicated and siControl. Blots were probed with different antibodies as indicated. γ-tubulin is used as a loading control. The normalized fold change in the expression with respect to control is indicated on the blots.

### Arl15 depletion alters HeLa cell migration and adhesion

Arl15 localizes to filopodia, and its knockdown in HeLa cells showed a reduced number of filopodia. Based on these observations, we hypothesized that loss of Arl15 possibly affects the migration of cells. We have performed a wound-healing assay at 0, 12, 24, and 48 h to quantify the differences in cell migration speeds. These results showed a significantly lower (0.39 folds) at 12 h, and a mild reduction at 24 or 48 h in cell migration of Arl15 depleted cells compared to control cells (**Fig. 7A**). To test the role of adhesion in migration, we suspended cells on fibronectin-coated coverslips for 30 and 60 min and stained cells with phalloidin Alexa Fluor 488. Cell spread areas were higher at both the time points (1.29 and 1.54 folds respectively) in siArl15 cells as compared to siControl cells (**Fig. 7B**). Interestingly, HeLa cell sizes, obtained using FACS were marginally higher (1.15 folds) upon Arl15 depletion compared to the control cells (**Supplementary Fig. 3E**). These studies showed that Arl15 knockdown cells display reduced cell migration and moderately enhanced cell adhesion and size.

**Figure 7.**
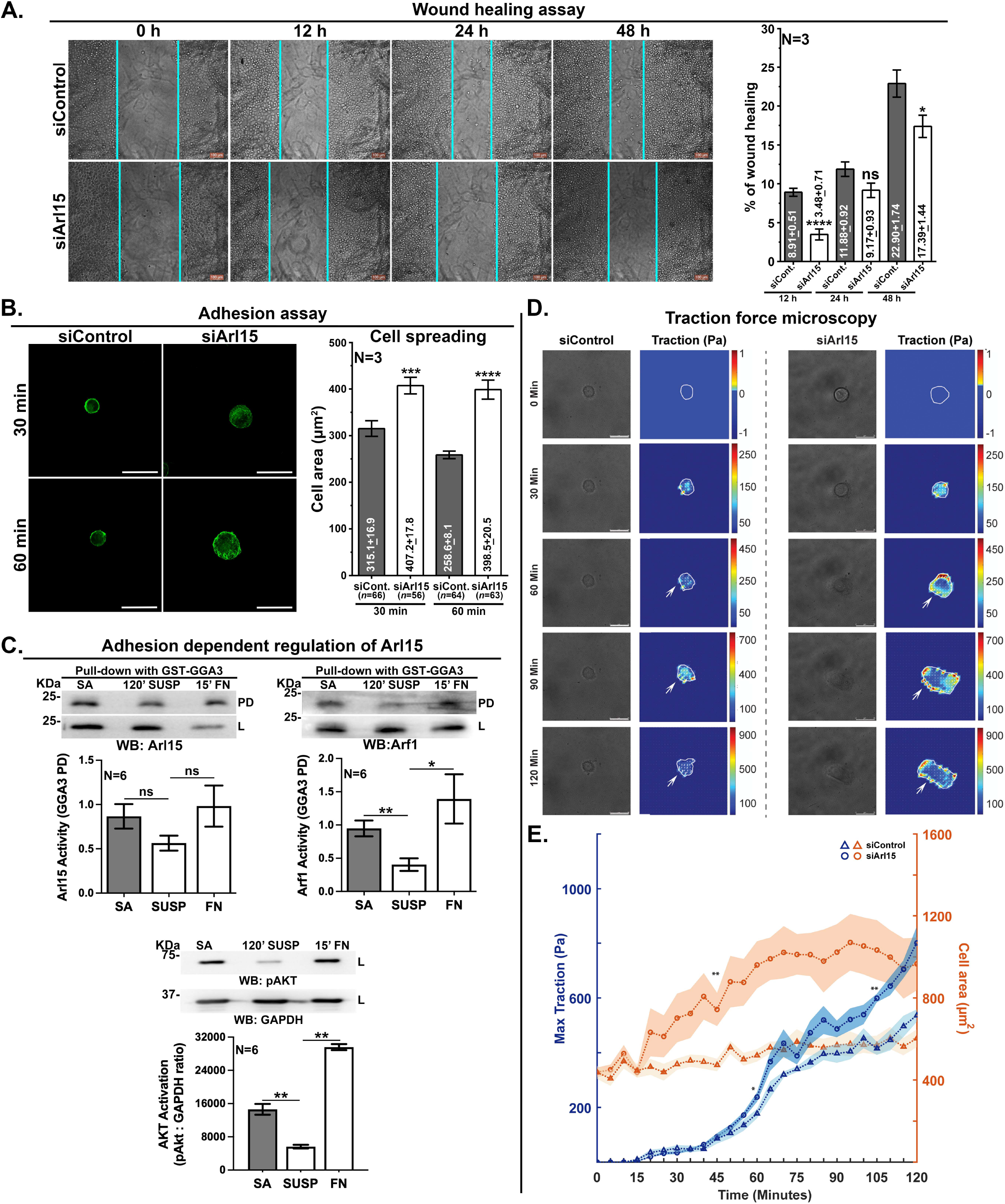
Arl15 depleted cells display reduced cell migration and enhanced cell spreading and adhesion. (A) siControl and siArl15 depleted HeLa cells were subjected to wound healing assay at different time points (0, 12, 24 and 48 h). The lines indicate the cell front used to measure the wound thickness. The graph represents the percentage of wound healing in siControl and siArl15 knockdown HeLa cells at different time points compared to 0 h. The average values in mean ± s.e.m. are indicated on the graph. Scale bars, 100 μm. N=3. (B) siControl and siArl15 knockdown HeLa cells were subjected to adhesion assay and measured the cell spread area at 30 min and 60 min of plating. Cells were fixed, stained with phalloidin Alexa Flour 488 and then quantified the cell area. Scale bar, 20 μm. The graph represents the measured cell areas (in μm^2^) at two different time points. The average values in mean ± s.e.m. are indicated on the graph. N=3, the number of repeats and n, the total number of cells. (C) Immunoblotting (WB) analysis of GST-GGA3 pull-down (PD) analytes from the cells grown on a tissue culture dish (stable adherent, SA), trypsinized and suspended in the medium for 120 min (120’ SUSP), and then subjected to adhesion for 15 min on fibronectin-coated (15’ FN) plate. GST-GGA3 pull-down (PD) from whole cell lysates (L) probed for Arl15 (left panel) and Arf1 (right panel) during the above conditions. The phosphorylation status of AKT (positive control, bottom panel) and total AKT levels in whole cell lysates (L) are shown separately. Graphs represent the average intensity of each pulldown band relative to whole cell lysate from six independent experiments. The graph representing the AKT activation was normalized with respective GAPDH values. N=6. (D) Time-lapse traction force microscopy of siControl and siArl15 HeLa cells. The image panel shows phase contrast images and the respective traction maps for the cells at 0, 30, 60, 90 and 120 min of the video. The traction force scale bar represents the constrained tractions exerted by the cells calculated using Fourier Transform Traction Cytometry (FTTC) method. The cell boundary from a phase-contrast image is used to calculate the tractions exerted by the cells. Scale bars, 25 μm. Note that the numbers of focal adhesion puncta (arrows) are higher for siArl15 cells than siControl cells at all time points. (E) The graph represents the average maximum traction (Pa) exerted by the six cells and their respective cell areas (μm^2^) at every five min of interval. The shaded region represents the standard deviation of maximum traction (orange) and cell areas (blue) for all six cells at each time point. \**P*≤0.05, \*\**P* ≤ 0.01, \*\*\**P* ≤ 0.001, \*\*\*\**P* ≤ 0.0001 and ns = non-significant.

Earlier studies showed that Arf family GTPases such as Arf1 (Singh et al., 2018) and Arf6 activation (Balasubramanian et al., 2007), which are implicated in intra-Golgi transport and endosomal recycling, are regulated by integrin-mediated cell adhesion (D’Souza-Schorey and Chavrier, 2006). Because Arf1 activation is required for Arl15 localization, we tested the role of integrin-mediated cell adhesion on the activation of Arl15 as described previously (Singh et al., 2018). We measured the activation of Arl15 in cells, serum-deprived but under stable adherent (SA) condition, which were detached and held in suspension for 120 min (120’ SUSP), and subsequently re-adhered on fibronectin (15’ FN) for 15 min. These experiments show that the activation of AKT (pS473), used as a positive control, dropped significantly during SUSP condition as compared to SA which was restored upon re-adhesion (15’ FN) (**Fig. 7C**) (Balasubramanian et al., 2007). We next measured the status of Arf1 activity based on the Golgi-localized γ-ear containing Arf-binding protein 3 (GGA3) pull-down during these conditions (Singh et al., 2018). Immunoblotting analysis of GST-GGA3 pull-down analytes showed reduced binding of Arf1 in 120’ SUSP compared to SA which was restored at 15’ FN condition similar to that of pAKT (**Fig. 7C**). Interestingly, GST-GGA3 pull-down of endogenous Arl15 from these lysates follows a kinetic pattern similar to Arf1 (**Fig. 7C**). However, the drop in levels of GST-GGA3 bound Arl15 at 120’ SUSP, restored at 15’ FN conditions were not significantly different from SA condition, unlike Arf1 (**Fig. 7C**). These results are consistent despite the pulldowns being repeated in six independent experiments. Thus, we predict that the activity of Arl15 may not be significantly affected by integrin-mediated cell adhesion like Arf1. This is further supported by the cell surface expression of β-integrin being unaffected by siArl15 compared to siControl cells (**Fig. 5A**).

To evaluate the role of Arl15 in adhesion, we used time-lapse traction force microscopy (TFM) using siControl and siArl15 HeLa cells on 10 kPa fluorescent bead coated polyacrylamide substrates as described earlier (Kulkarni et al., 2018). Phase-contrast images of Arl15 knockdown cells (n=5) showed increased spread areas compared to control cells (n=5) on polyacrylamide substrates (**Fig. 7D**). The numbers of focal adhesion puncta were also higher for siArl15 cells than siControl cells (**Fig. 7D**). FTTC tractions were lower during the initial adhesion phase and increased during the cell spreading phase. The maximum traction stresses exerted by the cell were 1.5 times higher for siArl15 cells (1067.73±78.30 Pa) as compared to siControl cells (714.17±29.56 Pa) at the end of 2 h after cell adhesion to the substrate (n=6 each group) (**Fig. 7E**). The total number of focal adhesion clusters were 1.44 times higher for siArl15 cells (7.67±0.67) as compared to control cells (5.33±0.61) at 120 min following cell seeding on the polyacrylamide gel substrates (**Fig. 7E**). The corresponding cell areas were 1.6 times higher for siArl15 cells (724.31±96.36 µm^2^) as compared to control cells (451.92±32.18 µm^2^) (**Fig. 7E**). The maximum tractions were statistically different (p <0.05) between the groups after 1 h of seeding, whereas the cell areas were statistically different (p <0.01) between groups after 45 min of seeding. Control cells took longer to spread than siArl15 cells **(Fig. 7D**). However, the cell areas were not statistically different between the groups with time at the end of 24 h post-seeding of cells on polyacrylamide substrates (data not shown). The maximum tractions were higher for the siArl15 (1451.86±111.78 Pa) group than siControl (544.11±61.73 Pa) 24 h after seeding the cells on the 10kPa polyacrylamide gel substrates (**Fig. 7E**). Overall, these results indicate that Arl15 plays a significant role in cell spreading and adhesion which has not been shown earlier.

## Discussion

The function of Arl15 in vesicular transport and its link to cellular processes has not been demonstrated. In this study, we characterized the intracellular function of Arl15 in HeLa cells. Earlier studies showed the localization of Arl15-GFP to Golgi and partly to the PM in differentiated adipocytes/C2C12 cells (Rocha et al., 2017; Zhao et al., 2017). Our extensive fluorescence microscopy analyses demonstrated the localization of Arl15-GFP predominantly to Golgi and PM along with filopodia and partly to REs in multiple mammalian cell types, including primary keratinocytes (**Fig. 1**). These data illustrate the conserved localization of Arl15 in many cell types. Intracellular localization of Arl15 was however independent of the actin cytoskeleton or the activity of Cdc42/Rac1/FAK1; these results suggest the direct association of Arl15 with the PM. The localization of Arl15 to Golgi however depends on the Golgi integrity or requires Arf1 activity. Loss of Arf1 expression did not affect Arl15 localization. These results implicate a compensatory mechanism for the Arf1 activity which may be mediated through other Arf GTPases in the Golgi (Pennauer et al., 2022; Volpicelli-Daley et al., 2005). Because a mutation at V80A in the GTP-binding domain blocks Arl15 localization to Golgi/PM, the GTPase cycle of Arl15 also contributed to its localization. Interestingly, V80A is a novel, non-classical dominant-negative mutation in Arl15 as compared to classical residues A86 or T46 in the GTP-binding domain, conserved in other Arls. Since none of the other mutations in Arl15 could make the protein cytosolic compared to Arl15^V80A^, our results indicate the uniqueness of this residue for Arl15 activity or required for the organelle localization. The predicted classical constitutive active (A86L) or myristoylation positive (S2G) mutations in Arl15 visually showed enhanced localization to Golgi/PM compared to Arl15^WT^. In contrast, the predicted dominant negative (T46N) mutation did not affect Arl15 localization (**Fig. 4**). *In silico* analysis of the Arl15 protein sequence by CSS-Palm predicted the three palmitoylation sites at the N-terminus (Cys17, Cys22, and Cys23); mutating the residues C22Y and/or C23Y abolished the localization of Arl15 to Golgi (Wu et al., 2021). In line to this study, we also predict that palmitoylation, but not myristoylation to play a key role in the recruitment of Arl15 to the microdomains at Golgi/PM (see below). Thus, our study has identified a novel amino acid, valine at the 80^th^ position (in the GTP binding domain) required for Arl15 function *in vivo*. Consistently, mutating the residue to alanine (V80A) significantly decreased the number of filopodia and moderately dispersed Golgi stacks in HeLa cells (**Fig. 4**).

Analysis of Arl15 depleted HeLa cells demonstrated that Arl15 is required for the trafficking of Golgi proteins such as caveolin-2 and STX6 to the cell surface (**Fig. 5**). Consistently, the depletion of Arl15 mislocalized these proteins to lysosomes and restored their localization to Golgi upon the treatment of cells with bafilomycin (**Supplementary Fig. 2**) but not with MG132 (data not shown). It is interesting to note that the SNARE STX6 has multiple functions which include (1) regulating the microdomain composition of PM by delivering the lipids from Golgi to support caveolar endocytosis (Choudhury et al., 2006); (2) retrograde transport of low-density lipoprotein (LDL) derived hydrolyzed cholesterol from the NPC1-positive endosomes to the TGN (Urano et al., 2008); and (3) regulating the delivery of acid hydrolases bound to MPR (mannose 6 phosphate receptor) from TGN to endosomes that are sorted by AP-1 (adaptor protein)-clathrin-coated buds during secretory granule maturation (Klumperman et al., 1998; Kuliawat et al., 1997). These studies indicate a role for STX6 in maintaining the organization of microdomains (positive for caveolae) in the PM. It is however unclear whether STX6 acts as a SNARE during these processes. Our study shows that Golgi localized Arl15 regulates STX6 dynamics and its associated cargo, such as caveolin-2 in the Golgi (Jung et al., 2012). The loss of caveolin-2 from Golgi may hence be due to the mislocalization of STX6 in Arl15 depleted cells, which requires future investigation.

Our study showed the dual localization of Arl15 majorly to both Golgi and PM. It is interesting to know whether the cell surface expressed Arl15 is regulated by the Golgi pool or vice versa. We predict that the Golgi pool of Arl15 regulates its cell surface localization due to the following observations: (1) treatment of cells with brefeldin A or golgicide A mislocalizes both pools of Arl15 to cytosol; (2) overexpression of Arf1^T31N^ results in membrane dissociation of both Arl15 pools; and (3) knockdown of Arl15 or expression of Arl15^V80A^ mutant in HeLa cells did not show any differential localization of Arl15. These results suggest that the Golgi pool maintains the PM localization of Arl15, and any disturbance in the Golgi pool affects its PM localization. In search of identifying the upstream factors for recruiting Arl15 to Golgi, we found interesting crosstalk with Arf1 but not with Arf family GEFs or GAPs based on the following results. The treatment of HeLa cells with Arf1-dependent Golgi dispersal agents such as brefeldin A/golgicide A or expression of the dominant-negative form of Arf1^T31N^ in HeLa cells mislocalize Arl15 to the cytosol, which reduces the number of filopodia. Thus, the activation of Arf1 or Arf1 GTPase cycle or Golgi integrity may be required to maintain the Golgi pool of Arl15 concurrently the PM pool (**Fig. 8**). Further, Arf1 or its homologues Arf3/4/5 in Golgi acts upstream of Arl15 for its recruitment to Golgi since the knockdown of Arf1 showed no effect on Arl15 localization. These data suggest the existence of compensatory mechanisms in the absence of Arf1 (Nakai et al., 2013; Pennauer et al., 2022; Volpicelli-Daley et al., 2005). However, Arf1 knockdown but not Arf family GEFs or GAPs reduces the expression of Arl15 in HeLa cells indicating a regulatory crosstalk between Arf1 and Arl15. We also observed a reduction in Arl15-dependent cargoes levels such as caveolin-2 in siGBF1 cells and SXT6 in siArf1 and siBIG2 cells (**Fig. 6**). These studies implicate the possible presence of other factors (addition to Arf1) in regulating the membrane recruitment of Arl15 at the Golgi. Studies have shown that ASAP1 acts as a GAP for Arf1 (Furman et al., 2002) and ASAP2 a proposed GAP for Arl15 (Zhao et al., 2017). Our extensive ASAP1/2 knockdown experiments in HeLa cells ruled out the possibility of these proteins acting as GAPs for Arl15. Nevertheless, the reason for the partial loss of STX6 and caveolin-2 localization from Golgi in the ASAP1/2 knockdown cells requires investigation. Overall, our data showed a crosstalk between Arl15 and Arf1 GTPase cycle.

**Figure 8.**
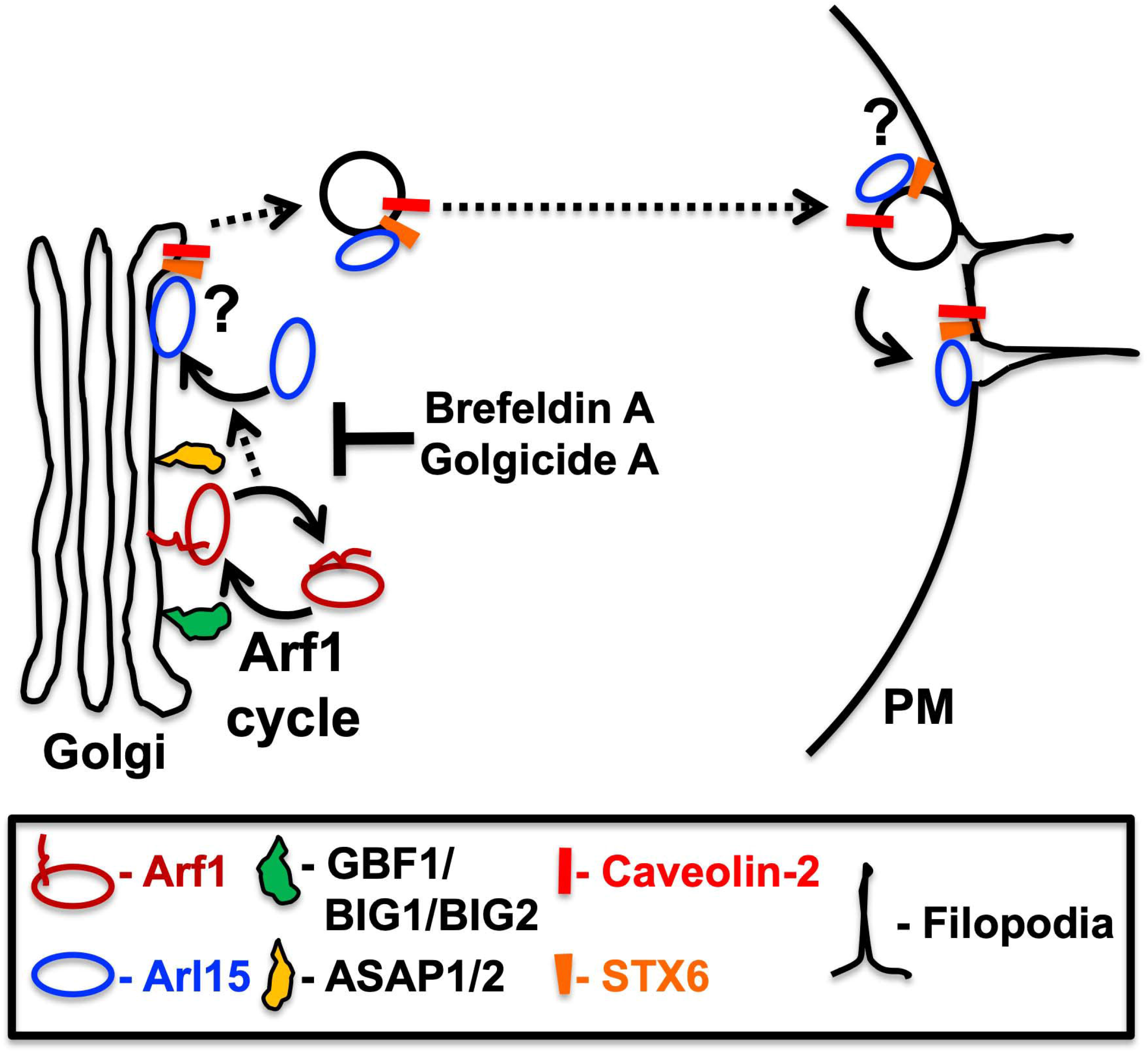
Proposed model for Arl15 function in vesicular transport and cell surface organization. Arl15 (Arl15-GFP) localizes majorly to Golgi and plasma membrane (PM), including filopodia. Our data suggest that the recruitment of Arl15 to Golgi is majorly dependent on the Arf1 GTPase cycle (indicated) or the Arf1-dependent Golgi integrity (T – shape, regulated by brefeldin A or golgicide A), but not directly by Arf family GEFs (GBF1, BIG1, BIG2) or GAPs (ASAP1, ASAP2). We predict that Arf1-GTP stabilizes the membrane association of Arl15 to Golgi (dashed arrow), which further regulates its localization to PM. The Golgi-associated Arl15 possibly regulates cargo sorting (?, requires investigation) and trafficking of several Golgi localized proteins such as caveolin-2 and STX6 to PM (dashed arrows). To facilitate this process, we hypothesized that Arl15 carries the cargo vesicle towards the PM to organize membrane domains composed of caveolin-2, STX6 etc. Moreover, these membrane domains possibly regulate the initial cell adhesion steps and may favour the biogenesis (length and number) of filopodia. The fusion of Arl15-positive vesicles with PM is possibly mediated by STX6 (?, requires investigation). As expected, the depletion of Arl15 mislocalizes the cargo and dysregulates the membrane domains at PM that possibly affecting the cell surface processes, including migration and adhesion.

Arl15 seems to regulate the PM organization at the cell surface. Our data in the Arl15 depleted cells showed: (1) decreased number of filopodia; (2) altered the vinculin organization to the focal adhesion sites; and (3) enhanced uptake of pinocytic and receptor-mediated endocytic cargo without affecting the cargo recycling (**Fig. 5**). These changes are likely made feasible due to alteration in PM organization, especially the microdomains that are positive for caveolin-2 and STX6 proteins. Based on the results, we propose a model such that: (1) Arf1 GTPase cycle recruits Arl15 to the Golgi; (2) at the TGN, Arl15 regulates the sorting of caveolin-2, STX6 and other cargo into vesicles/tubules (REs) by an as yet unknown mechanism; (3) the Arl15 positive vesicles/tubules are targeted and fused with PM possibly through STX6 (indicated with ‘?’) and deliver the cargo (**Fig. 8**). This process possibly regulates the microdomain organization at the PM, which is important for cell spreading, adhesion, and filopodial formation.

The proposed model for the function of Arl15 supports our results: first, we show reduced filopodial numbers in siArl15 and Arl15^V80A,A86L,E122K^ mutant expressing cells. Second, cargo trafficking (pinocytosis and receptor-mediated endocytosis) is altered at the PM in siArl15 cells. Finally, we suggest a key function for Arl15 at the PM which is based on reduced migration, enhanced cell spreading and increased number of focal adhesion points with increased tractions (at 2 h of adhesion) in siArl15 cells. The binding of Arl15 to GGA3 and mislocalization of cargo at Golgi in siArl15 cells suggest that Arl15 also functions at TGN in contrast to other functions such as modulation of magnesium transport in Golgi, reported recently (Zolotarov et al., 2021). We also noticed the localization of Arl15 to tunnel-like nanotubes (not shown) between the Neuro 2a cells suggesting a possible role for Arl15 in cell-cell communication. Nevertheless, Arl15 possibly plays a key role in regulating filopodial biogenesis (length and number) or may control the trafficking of important cargoes to its site of origin, which needs further investigation. Vinculin is enriched in integrin-based focal adhesion points that connect the extracellular matrix (ECM) (Atherton et al., 2016). In 2-dimensional (2D) cell culture models, vinculin inhibits cell migration by stabilizing cell adhesions (Mierke et al., 2010). In contrast, vinculin promotes directionally persistent cell migration by generating the traction force required for cell motility and ECM remodeling in 3D collagen (Thievessen et al., 2015). Our data show the enhanced cell spread and adhesion in siArl15 cells, which may be due to a change in vinculin assembly at the focal adhesion sites. Overall, these studies highlighted the molecular function of Arl15 in protein sorting from the Golgi to the PM to modulate various cellular processes, including filopodia formation. We also identified the molecular players that regulate the membrane recruitment of Arl15 to Golgi which control its PM localization/function. Further, the identification of Arl15 effector molecule/s that possibly control Arl15-associated cellular processes and metabolic traits requires future investigation.

## Materials and Methods

### Reagents and antibodies

All chemicals and tissue culture reagents were purchased from Sigma-Aldrich (Merck) or ThermoFisher Scientific (Invitrogen). CAS 1177865-17-6 (RAC1 inhibitor, 553502), ML 141 (CDC42 inhibitor, SML0407), PF-573228 (FAK inhibitor, PZ0117), bafilomycin A1 (B1793), brefeldin A (B7651), fibronectin human plasma (F2006), Golgicide A (G0923), latrunculin A (L5163), methyl cellulose (M0262), MG132 (C2211) and nocodazole (M1404) were procured from Sigma-Aldrich. Dextran Fluorescein (70K MW, D1822), Phalloidin-Alexa Fluor 488 (A12379), Phalloidin-Alexa Fluor 594 (A12381), Transferrin (human)-Alexa Fluor 594 (T13343) and Wheat Germ Agglutinin (WGA)-Alexa Fluor 594 (W11262) were obtained from ThermoFisher Scientific (Invitrogen). Matrigel was purchased from BD Biosciences.

The following commercial polyclonal and monoclonal antisera were used (m, mouse; h, human and r, rat proteins). Abcam: anti-mArf1 (ab2806 for IP) and anti-hArf1+Arf3 (ab32524). BD Biosciences: anti-rGM130 (610822) and anti-hp230/golgin-245 (611281). Developmental Studies Hybridoma Bank: anti-hLAMP-1 (H4A3). Cell Signaling Technology: anti-mAkt (9272), anti-Phospho-mAkt (Ser473) (9271), anti-hCalnexin (2679), anti-hCaveolin-1 (3267), anti-hCaveolin-2 (8522), anti-hEEA1 (3288), anti-hLAMP1 (9091) and anti-mSyntaxin 6 (2869). Invitrogen: anti-hArf1/Arf3/Arf5/Arf6 (MA3-060), anti-hEGFR (MA5-13319) and anti-GFP (A-11122). Proteintech: anti-hArl15 (11934-1-AP). Santacruz biotechnology: anti-hDDEF1 (ASAP1, sc-374410), anti-hDDEF2 (ASAP2, sc-374323), anti-hBIG1 (sc-376866), anti-hBIG2 (sc-398042), anti-GAPDH (sc-25778), anti-hGBF1 (sc-136240), anti-hGRASP65 (sc-365434) and anti-hIntegrin β1/ITGB1 (sc-13590). Sigma-Aldrich: Anti-mGAPDH (G9545), anti-Vinculin (V4505) and anti-γ-Tubulin (T6557). All secondary antibodies were either from Invitrogen or Jackson Immunoresearch.

### siRNAs and plasmids

#### siRNAs

The following target siRNA sequence against the respective gene were synthesized from Eurogentec, Belgium. hArl15 (5’-GCUUCUCUCAGCUGAUUAA-3’), hArl15-3’-UTR (5’-CUCACACUAUAUUACAGAA-3’), hArf1 (5’-UUCCCACCAUAGGCUUCAA-3’), hGBF1 (5’-CCUCUGUCAACAAGUUCCU-3’), hBIG1 (5’-GGAAUCUGGAAAUUCUUCA-3’), hBIG2 (5’-CGACUGUGAUUUAAAUGCU-3’), hASAP1 (5’-GACCUGACAAAAGCCAUUA-3’) and hASAP2 (5’-AGAGGACUCCCAAAUUCGU-3’).

#### Plasmids

mArl15^WT^-pEGFP-N3 construct was made by cloning the PCR amplified Arl15 from mouse cDNA prepared from mouse melanocytes (melan-Ink4a-Arf-1). The Arl15 PCR product was digested with *Bgl*II and *Xho*I restriction enzymes, followed by cloning into pEGFP-N3 at *Bgl*II and *Sal*I sites. Similarly, mArl15^S2G^-pEGFP-N3 was prepared by amplifying the Arl15 gene with a forward primer having S2G mutation. The PCR product was digested with *Bgl*II and *Xho*I enzymes and then subcloned into pEGFP-N3 at *Bgl*II and *Sal*I sites. The constructs were confirmed by restriction digestion and DNA sequencing. Site-directed mutagenesis of Arl15^WT^-GFP or Arl15^S2G^-GFP was carried out using a protocol described previously (Edelheit et al., 2009) and generated Arl15^S2A^-GFP, Arl15^T46N^-GFP, Arl15^A86L^-GFP and Arl15^A86Q^-GFP (shown in Figure 4). All constructs were sequenced before use. Unfortunately, we found additional mutations at V80A, E122K in Arl15^A86L^-GFP during sequencing analysis. To confirm the role of these additional mutations, we have generated individual and combination mutations, as shown in Figure 4. mArl15^WT^-GFP-pLVX-puro construct was prepared by subcloning the *Bgl*II and *Xba*I fragment of Arl15-GFP from pEGFP-N3 into *Bam*HI and *Xba*I sites of pLVX-puro vector (Clontech), prepared in *dam*^*-*^*dcm*^*-*^ *E. coli* strain.

Arf1^WT^-GFP, Arf1^T31N^-GFP and Arf1^WT^-mCherry are obtained (Kumari and Mayor, 2008) as a gift from Prof. Jitu Mayor, NCBS, Bangalore, India. pMD2.G (12259), psPAX2 (12260), mCherry-UtrCH (26740) were obtained from Addgene. KIF13A-YFP has been described (Delevoye et al., 2009).

### Protein homology modelling

A BLAST search of ARL15 identified its closest homologues as ARL2 (UniProt - P36404) and ARF6 (UniProt - P62330), which each share ∼34% sequence identity with ARL15. The crystal structure of ARL2 in complex with BART (PDB ID: 3DOE) was chosen as a template and built the Arl15^WT^ models using iTasser server (Roy et al., 2010; Yang et al., 2015; Yang and Zhang, 2015). Further, the iTasser models were refined using Galaxy Refine (Ko et al., 2012). The final models were sourced from the AlphaFold2 protein structure database, and the mutants were modelled using AlphaFold Colab (Jumper et al., 2021). The models were analysed in PyMOL and surface electrostatics using APBS module (Baker, 2004).

### Tissue culture, transfection and lentivirus viral transduction

HeLa, HEK293T, Neuro 2a (N2a) and A549 cell lines (from ATCC) were maintained in complete DMEM medium [DMEM (Invitrogen)+10% FBS (Biowest)+1% Pen-Strep (Invitrogen) + 1% L-glutamine (Invitrogen)]. Cells were grown at 37ºC in a humified cell culture chamber supplied with 10% CO_2_. Neonatal human primary epidermal keratinocytes (NHEK) were purchased from Lonza, maintained in EpiLife Serum-free medium supplemented with HKGS (Invitrogen) and incubated at 37ºC. Arl15-GFP stably expressing HeLa cells were also maintained in a complete DMEM medium supplemented with puromycin (1 µg/ml). For the transfection of the plasmids, mammalian cells in a 35 mm dish were starved for 30 min in OPTI-MEM (Invitrogen) medium, followed by the addition of a mix made with DNA (1 µg) and Lipofectamine 2000 (3µl; Invitrogen). Cells were changed with complete DMEM post 6 h and then incubated for 48 h at 37ºC. Finally, cells were fixed, stained and imaged. For siRNA-mediated knockdown, HeLa cells were seeded at 60 - 70% confluency and added with a premix containing 120 nM of siRNA and 5 µl of oligofectamine made in OPTI-MEM medium. Post 8 - 10 h of transfection, cells were replaced with complete DMEM and then incubated at 37ºC for a total of 48 - 52 h of transfection before use. For immunoblotting, cells were harvested between 48 - 52 h of transfection. For immunofluorescence microscopy, cells were plated on coverslips post 24 h of transfection, fixed after 48 h, stained and imaged.

Stable cells expressing Arl15-GFP in HeLa cells were prepared by lentivirus transduction, followed by gradually selecting the cells with puromycin (1 - 5 μg/ml) for 15 days. A non-clonal population of cells (referred to here as HeLa:Arl15-GFP stable cells) was used for the experiments. The cells were maintained in media containing 1 µg/ml puromycin. Lentivirus using HEK293T cells is prepared following a previously described protocol (Shakya et al., 2018).

### Measurement of relative cell size using flow cytometry

HeLa (plain) and HeLa:Arl15-GFP stable cells were seeded at a 50% confluency in a complete DMEM medium for 24 h without and with puromycin (1 µg/ml), respectively. The cells were trypsinized, washed and resuspended in 300 µl 1X ice-cold PBS. The samples were loaded on FACSVerse (BD Biosciences) system and gated for the GFP channel. The intensity values for a minimum of 10,000 cells were acquired per sample and analyzed using FlowJo software (BD Biosciences). The median values of forward scatter (represents the relative size) were taken to calculate the fold change between HeLa (plain) and HeLa:Arl15-GFP stable cells and then plotted. Similarly, the fold change in relative size between siControl and siArl15 cells was calculated and then plotted.

### Internalization of fluorescein-conjugated dextran and recycling kinetics of transferrin (Tf)-Alexa Fluor 594

For fluorescein-conjugated dextran internalization, both siControl and siArl15 HeLa cells were incubated on ice with 0.5 mg/ml of fluorescein-dextran beads (70,000 MW) for 30 min and then continued the incubation for 2 h at 37ºC in a CO_2_ incubator. Post chase, cells were washed with 1X PBS, fixed with 3% formaldehyde and then imaged. The intensity of intracellular accumulated fluorescein-dextran was calculated by measuring the corrected total cell fluorescence (CTCF) using Fiji software and then plotted.

Internalization and recycling of Tf-Alexa Fluor 594 in siControl siArl15 HeLa cells were performed as described previously (Shakya et al., 2018). Briefly, cells on coverslips were starved in a serum-free medium containing 25 mM HEPES for 30 min at 37ºC. Next, cells were transferred and incubated on ice for 30 min in a complete medium (buffered with 25 mM HEPES) containing Tf-Alexa Fluor 594 (30 μg/ml) (referred to here as binding on ice). A set of coverslips were transferred to a CO_2_ incubator at 37ºC for 20 min (referred to here as 20 min uptake). Another set of coverslips was washed with 1X PBS, transferred to plain complete medium and then chased for 40 min at 37ºC in a CO_2_ incubator (referred to here as 40 min chase). At the end of each step, the cells were fixed with 3% formaldehyde and imaged. Images were captured under identical camera settings in a fluorescence microscope. The fluorescence intensity in each condition was measured as CTCF using Fiji and then plotted.

### Cell surface binding of wheat germ agglutinin-Alexa Fluor 594 (WGA-AF594)

Cells in suspension were washed and incubated with WGA-AF594 (0.5 ng/µl) in 200 µl of HBSS buffer for 15 min on ice in dark condition. Cells were washed with 1X PBS and then fixed with 3% formaldehyde for 15 min at room temperature. Cells were washed and suspended in 200 µl of 1X PBS and analyzed by BD CytoFLEX LX cell analyzer. The mean fluorescence intensity (MFI) for each sample was calculated and then plotted.

### Staining of cells and immunofluorescence microscopy

Cells on coverslips were fixed with 3% formaldehyde for 15 - 20 min at room temperature and then washed and stored in 1X PBS with 0.05% sodium azide until further use. Cells were stained with primary antibodies, washed with 1X PBS and then stained with respective secondary antibodies as described previously (Jani et al., 2015). In a few experiments, cells on coverslips were subjected to internalization of Tf-Alexa Fluor 594 or fluorescein-dextran, followed by fixation of cells with 3% formaldehyde. In some experiments, cells were stained with phalloidin-Alex Fluor 488/594 (2 U/ml) in 1X PBS or Wheat Germ Agglutinin (WGA)-Alexa Fluor 594 (50 μg/ml) in HBSS for 30 - 60 min. Immunofluorescence microscopy (IFM) of cells was carried out using the microscope setup described previously (Jani et al., 2015) or Zeiss LSM 780 with Airyscan module (for super-resolution imaging).

### Quantification methods of IFM

#### 1. Quantification of fluorescence intensity by corrected total cell fluorescence (CTCF)

Merge the image layers into single MIP (maximum intensity projection) image using ImageJ (Image tab >> stacks >> z-projection > set the range of z-stacks that are supposed to be merged >> OK >> separate MIP image will be rebuild). Measure the following parameters required for CTCF calculation: area of the cell and mean fluorescence and integrated density (Go to Analyze >> set measurement >> select Area, Mean grey value and integrated density >> OK). Use the freehand selection tool to select the individual cell in the field and then note the parameters (Go to Analyze >> Measure >> gives result separate result sheet >> copy the results to Excel). Similarly, the background intensity was calculated for each quantified image. The CTCF value for each cell = Integrated density – (Area of the cell * Mean fluorescence of the background). The average CTCF for each set was calculated and then plotted.

#### 2. Quantification of the number of fluorescent puncta

Open the IFM images in ImageJ and select the single channel using the split channel tool present under the image tab within color dropdown. Merge the z-stacks into a single plane using the z-projection tool present within the stacks dropdown under the image tab. Convert the selected single-channel MIP image into 8 bit followed by rolling ball background subtraction (Process tab >> Subtract Background) with 20.0 pixels radius. Further, adjust the image to auto thresholding (30 to 255 as default; Image tab>>Adjust>>Threshold). Next, convert the image to binary (Process tab>>Binary>>Make binary), succeeding with watershed segmentation (Process tab>>Binary>>Watershed). Calculate the number of puncta within a relative cell area using the freehand selection tool (Analyze tab>>Analyze particles). The obtained punctum counts are relatively based on default values of particle size (zero to infinity μ^2^) and circularity (0.00-1.00). The average punctate number per cell was calculated and then plotted.

#### 3. Quantification of the percentage of cells with caveolin-2 staining

HeLa cells were stained with anti-caveolin-2 and anti-GM130 (to label Golgi) antibodies and then imaged. The cells with Golgi localized caveolin-2 staining were visually counted to a total number of cells within the field. The percentage of cells with caveolin-2 staining was calculated and then plotted.

### Isolation of RNA and preparation of cDNA

Total RNA from HeLa cells (human) or melanocytes (mouse) melan-Ink4a (Shakya et al., 2018) was isolated using a GeneJET RNA purification kit (ThermoFisher Scientific) in the presence of β-mercaptoethanol. The RNA concentration was estimated using a NanoDrop 2000C spectrophotometer (ThermoFisher Scientific). The cDNA was prepared from total RNA using the RevertAid First Strand cDNA synthesis kit (K1622, ThermoFisher Scientific). The human or mouse Arl15 was amplified from respective cDNA using specific primers.

### Immunoblotting

The harvested cell pellets were lysed in RIPA buffer and subjected to immunoblotting as described previously (Shakya et al., 2018). Protein estimation was carried out using Bradford reagent (Bio-Rad), and an equal amount of lysates was used for SDS-PAGE electrophoresis. Immunoblots were developed with the Clarity Western ECL substrate (Bio-Rad), imaged in a Molecular Imager ChemiDoc XRS+ imaging system (Bio-Rad) and analyzed using Image Lab 4.1 software. Protein band intensities were measured using Image Lab software and then normalized with γ-Tubulin. The fold change in protein band intensities with respect to control was calculated and indicated in the figure.

### Measurement of cell migration by wound healing assay

Cells were plated in duplicated wells of 24 well plate (0.15×10^6^ cells/well) to form a compact monolayer. Post 12 h, two different wounds were made by scratching the cells perpendicularly to the plate using a toothpick. Cells were washed twice with 1X PBS and then supplemented with serum-free DMEM medium, followed by imaging of the wound area (considered as 0 h). Similarly, images were taken every 12 h of incubation at 37ºC until the 48 h time point. The scratch distance was measured at three different positions per image, and then calculate the average distance at different time points 0, 12, 24 and 48 h. The observed distance at 0 h is considered the initial distance. The % of wound healing was calculated using the formula: % of wound healing = (initial distance-final distance/initial distance)*100. The average % wound healing at different time points was calculated and then plotted.

### Measurement of cell spreading by adhesion assay

HeLa cells were incubated in 0.2% serum containing medium for 12 h. Post incubation, cells were trypsinized, washed and reseeded onto overnight coated fibronectin coverslips. Cells were fixed at 30 and 60 min of adhesion with 3% formaldehyde. Further, cells were stained with phalloidin-Alexa Fluor 488 and then imaged in a confocal microscope with a 40X objective. The cell area (based on phalloidin-Alexa Fluor 488 fluorescence) was calculated using the freehand selection tool in ImageJ and plotted the average cell area at different timepoints.

### Measurement of adhesion-dependent regulation of Arl15 by GGA3-pulldown assay

Mouse embryonic fibroblasts (WT-MEFs, kind gift from Dr Richard Anderson, University of Texas Health Science Center, Dallas, TX) were cultured in DMEM supplemented with 5% FBS and 1% Pen-Strep at 37ºC in a 5% CO_2_ incubator. WT-MEFs (10^6^) were serum-deprived for 12 h in low-serum DMEM, detached with trypsin-EDTA and kept in suspension with 1% methylcellulose for 120 min (referred to here as 120′ SUSP). Further, cells were re-plated on fibronectin (10 µg/ml) coated plates for 15 min (referred to here as 15′ FN) or for 4 h to become stable adherent (referred to here as SA). At the end of each time point, cells were frozen, lysed in activity assay buffer, and then measured the activated Arf1 and Arl15 by pulldown of GST-tagged GGA3 (Golgi-localized γ-ear containing Arf-binding protein 3, known to bound to Arf1) as described earlier (Pawar et al., 2016). The cell lysate equivalent to 400 µl was used for pulldown assay and then eluted in 20 µl of Laemmli buffer. The samples were subjected to 12.5% SDS-PAGE by loading 2.8% of the lysate (22.5 µl of 400 μl) and a total 100% eluent of GST-GGA3 pulldown assay. Immunoblotting was performed and then probed with respective antibodies, as detailed in **Fig. 7C**. Blots were developed with the PICO chemiluminescence detection system (ThermoFisher Scientific) and then imaged in ImageQuant LAS 4000 (Fujifilm-GE). The band intensities were quantified using Image Lab software.

### Measurement of force during the cell adhesion by traction force microscopy

Polyacrylamide (PA) gels of 10 kPa stiffness were prepared as described earlier (Kulkarni et al., 2018). Briefly, 3-amino-propyl triethoxy silane (Sigma-Aldrich) treated coverslips were prepared, incubated at room temperature for 15 min, and then washed thoroughly in double-distilled water. Coverslips were next incubated in 0.5% glutaraldehyde solution (S D Fine-Chem Ltd.) for 30 min and air-dried. Another set of clean coverslips was treated with poly-L-lysine solution (Sigma-Aldrich) for 45 min at 37ºC in a humid chamber to activate the surface. The fluorescent bead solution (18 µl/ml) was spin-coated on the activated coverslip at 800 rpm for 5 min. Solutions of acrylamide (40% wt/vol, Sigma-Aldrich) and N, N’-methylene bisacrylamide (2% wt/vol, Sigma-Aldrich) were mixed with distilled water to obtain gels of 10 kPa stiffness (Tse and Engler, 2010). A solution of 30 μl of the PA gel with 1/100 total volume of APS (ammonium persulfate) and 1/1000 total volume of TEMED was sandwiched between one clean coverslip and another bead coated one at room temperature for 30 min. The bead-coated coverslip was carefully removed, and the PA gel-coverslip was attached to a 35 mm punched petri dish (Nunc, ThermoFisher Scientific) using a thin layer of vacuum grease (S D Fine-Chem Ltd.). A stock solution (100 mg/ml) of heterobifunctional sulpho-SANPAH linker (Pierce, ThermoFisher Scientific) was diluted to 1:200 in 50 mM HEPES buffer (Sigma-Aldrich) and then pipetted to the gel surface. Finally, the assembly was exposed to 365 nm UV light (ThermoFisher Scientific) for 10 min. Fibronectin (40 μg/ml, Sigma-Aldrich) was added to the gel surface and incubated at 37ºC for 45 min in a humid chamber.

The assembly was rinsed thrice with HEPES buffer, and the cells were seeded at 2000 cells/ml (total volume 2 ml) before starting the experiment at 37ºC supplemented with 5% CO_2_ in a live cell chamber (Leica DMI6000B with PEKON stage). Cells that attached and spread during the first two hours were used to quantify the changes in traction stresses during adhesion. Three images were obtained for each cell using a 40X oil immersion objective: (1) a phase-contrast image of the cell; (2) a fluorescent image of beads on the gel surface with attached cell (stressed configuration); and (3) a fluorescent image of beads (referential configuration) obtained following removal of the cell from the gel substrate using trypsin (Sigma-Aldrich).

Images of the cell and bead positions in the stressed and referential configurations were used to quantify the (constrained) traction stresses exerted by cells on the substrate using the Fourier Transform Traction Cytometry method in MATLAB (Kulkarni et al., 2018). Image drift correction was performed using Fiji by ImageJ. Experiments were performed three times for each set of the siControl and siArl15 cells (n=6 in each group). Data are reported as mean ± SEM. Results from the different groups were compared using a one-way analysis of variance (ANOVA) with Bonferroni comparisons to test for possible differences at each time point and indicate the significant difference on the graph. Cell tractions were also quantified 24 h after attachment to the 10 kPa polyacrylamide substrates using the same method (n=10 in each group).

### Statistical analysis

All statistical analyses were carried out using GraphPad Prism 6.04. All experiments are performed as experimental triplicates, and cell numbers are indicated in the graphs. The statistical significance between the groups was estimated by an unpaired Student’s *t*-test. Mann–Whitney’s test was used to calculate the significance of the adhesion-dependent regulation of the Arl15 assay. \**P*≤0.05, \*\**P* ≤ 0.01, \*\*\**P* ≤ 0.001, \*\*\*\**P* ≤ 0.0001 and ns = non-significant.

## Acknowledgement

We thank Jitu Mayor (NCBS) for the generous gift of the Arf1 constructs. We acknowledge Sachin Kotak (IISc) for sharing plasmids and reagents. We thank V Niranjan Kumar for the purification and pulldown experiments with His-Arl15 (data not shown) and Kavya Gopal for cloning the human Arl15. We acknowledge L. Alfonso Martinez-Cruz and Michel L. Tremblay for sharing the Arl15 coordinates (Zolotarov et al., 2021).

## Author contributions

PS performed all the experiments in this study. PHV supported the study with the siRNA-mediated knockdown experiments and quantification of IFM and FACS experiments. NP and NG conducted the traction force microscopy experiments. NS, RBR and NB performed the adhesion-dependent Arf/Arl activation experiments. AD and AP modelled the Arl15 structure and analyzed the mutations generated in this study. SRGS designed, oversaw the entire project, coordinated and discussed the work with co-authors and wrote the manuscript.

## Supplementary information

**Supplementary Figure 1. Mouse Arl15 is homologous to human Arl15 and localizes to Golgi but not to other intracellular compartments**. (A) The predicted structural view of human and mouse Arl15 were generated through homology modelling. The comparison between human and mouse (m/h) sequences is shown separately. The numbers in the comparison structures represent the amino acids of the mouse, which differ from the human sequence. (B) Immunoblotting analysis of plain, GFP-expressing and Arl15-GFP stably expressing HeLa cells. The blots were probed with anti-GFP and anti-GAPDH (loading control) antibodies. HeLa:Arl15-GFP stable cells showed a cohort of GFP cleavage in the cell lysate. (C) IFM analysis of HeLa:Arl15-GFP stable cells. Cells were stained for p230/golgin-245 (Golgi), calnexin (ER), EEA1 (early endosomes), LAMP-1 (lysosomes) and vinculin (focal adhesion) separately. Insets are magnified views of the white boxed areas. Scale bars, 10 µm. (D) Plot represents the fold change in cell size between plain HeLa and HeLa:Arl15-GFP stable cells (HeLa:A15G) by flow cytometry analysis. The relative fold change in values (mean ± s.e.m.) are indicated on the graph. N=3. ns = non-significant.

**Supplementary Figure 2. (A) Arf1 depletion reduces the expression of Arl15**. Immunoblotting analysis of control, Arf1 depleted, and Arl15 depleted HeLa cells for the expression of Arf1 and Arl15. γ-tubulin is used as a loading control. *, the non-specific band developed with the antibody. The normalized fold change in the expression with respect to control is indicated on the blots. **(B) 3’-UTR specific Arl15 siRNA reduces the Arl15 expression equivalent to cDNA specific siRNA**. Immunoblotting analysis of control and Arl15 knockdown (with gene or 3’-UTR specific siRNAs) HeLa cells. γ-tubulin and GAPDH are used as loading controls. *, the non-specific band developed with the antibody. **(C) Arl15 depletion does not change the Arf1 localization to Golgi**. IFM analysis of siControl and siArl15 treated HeLa cells. Cells were transfected with Arf1-GFP and then stained with an anti-GM130 antibody. Arrows point to Golgi localized Arf1-GFP and is moderately dispersed in siArl15 compared to siControl. In all images, insets are magnified views of the white boxed areas. Scale bars, 10 µm. **(D, E) Arl15 depletion mislocalizes the caveolin-2 and STX6 but not caveolin-1, EGFR or LAMP-1. Treatment of Arl15 knockdown cells with bafilomycin A1 restored the localization of caveolin-2 and STX6 to Golgi**. IFM analysis of siControl and siArl15 treated HeLa cells. Cells were stained for caveolin-1, LAMP-1 or EGFR (in D). In E, cells were treated with or without bafilomycin A1 and then stained with caveolin-2 or STX6 and LAMP-1. Arrows point to the mislocalization of caveolin-2 or STX6 in siArl15 cells and restored upon treatment with bafilomycin A1. Insets are magnified views of the white boxed areas. Scale bars, 10 µm. (F) Immunoblotting analysis of siControl and siArl15 treated HeLa cells. Cells were treated with or without bafilomycin A1 or MG132. Blots were probed for caveolin-2 and STX6 proteins. γ-tubulin is used as a loading control. The normalized fold change in the expression with respect to control is indicated on the blots. (G) **Arl15 knockdown cells accumulate the internalized transferrin and display no effect on its recycling**. IFM analysis of siControl and siArl15 knockdown HeLa cells incubated with transferrin Alexa Fluor 594 on ice (binding on ice), then incubated at 37ºC for 20 min (uptake) and washed and then chased with plain medium for 40 min (chase). Insets are magnified views of the white boxed areas. Scale bars, 10 µm. (H) Plot represents the intensity (CTCF) of bound/internalized/recycled transferrin Alexa Fluor 594 (F) in siControl and siArl15 treated HeLa cells. A.U., arbitrary units. The average values in mean ± s.e.m. are indicated on the graph. N, the number of repeats and n, the total number of cells. (I) **Arl15 depletion alters the lectin cell surface expression in HeLa cells**. The plot represents the fold change in lectin (WGA-Alexa Fluor 594) cell surface expression between siControl and siArl15 HeLa cells by flow cytometry analysis. The relative fold change in values (mean ± s.e.m.) are indicated on the graph. N=9. \*\**p*≤0.01 and \*\*\*\**P* ≤ 0.0001.

**Supplementary Figure 3. Validation of siRNA-mediated knockdown and the localization of GBF1 in siArf1 or Arf1-GEFs/GAPs depleted HeLa cells**. (A) Immunoblotting analysis of siControl, siArf1, siArf family GEFs (GBF1, BIG1, BIG2) and siArf family GAPs (ASAP1, ASAP2) treated HeLa cells. Blots were probed with respective antibodies as indicated. γ-tubulin is used as a loading control. *, the non-specific band developed with the antibody. (B) IFM analysis of pooled siRNA mediated knockdown in HeLa:Arl15-GFP stable cells as indicated. (C) IFM analysis of siRNA mediated knockdown of HeLa cells as indicated. Cells were stained with anti-GBF1 antibody. In all IFM images, insets are magnified views of the white boxed areas. Scale bars, 10 µm. (D) Plots represent the intensity (CTCF) of caveolin-2 or STX6 in siControl, siASAP1 (siA1) and siASAP2 (siA2) treated HeLa cells. A.U., arbitrary units. The average values in mean ± s.e.m. are indicated on the graph. N=3, the number of repeats and n, the total number of cells. (E) Plot represents the fold change in cell size between siControl and siArl15 HeLa cells by flow cytometry analysis. The relative fold change in values (mean ± s.e.m.) are indicated on the graph. N=3. \**P*≤0.05, \*\**P* ≤ 0.01, \*\*\*\**P* ≤ 0.0001 and ns = non-significant.

## References

Akisik, E., H. Yazici, and N. Dalay. 2011. ARLTS1, MDM2 and RAD51 gene variations are associated with familial breast cancer. Mol Biol Rep, 38:343–348.

Arya, S.B., G. Kumar, H. Kaur, A. Kaur, and A. Tuli. 2018. ARL11 regulates lipopolysaccharide-stimulated macrophage activation by promoting mitogen-activated protein kinase (MAPK) signaling. The Journal of biological chemistry, 293:9892–9909.

Atherton, P., B. Stutchbury, D. Jethwa, and C. Ballestrem. 2016. Mechanosensitive components of integrin adhesions: Role of vinculin. Experimental cell research, 343:21–27.

Baker, N.A. 2004. Poisson-Boltzmann methods for biomolecular electrostatics. Methods Enzymol, 383:94–118.

Balasubramanian, N., D.W. Scott, J.D. Castle, J.E. Casanova, and M.A. Schwartz. 2007. Arf6 and microtubules in adhesion-dependent trafficking of lipid rafts. Nature cell biology, 9:1381–1391.

Barral, D.C., S. Garg, C. Casalou, G.F. Watts, J.L. Sandoval, J.S. Ramalho, V.W. Hsu, and M.B. Brenner. 2012. Arl13b regulates endocytic recycling traffic. Proc Natl Acad Sci U S A, 109:21354–21359.

Behnia, R., B. Panic, J.R. Whyte, and S. Munro. 2004. Targeting of the Arf-like GTPase Arl3p to the Golgi requires N-terminal acetylation and the membrane protein Sys1p. Nature cell biology, 6:405–413.

Bock, J.B., J. Klumperman, S. Davanger, and R.H. Scheller. 1997. Syntaxin 6 functions in trans-Golgi network vesicle trafficking. Molecular biology of the cell, 8:1261–1271.

Burd, C.G., T.I. Strochlic, and S.R. Setty. 2004. Arf-like GTPases: not so Arf-like after all. Trends in cell biology, 14:687–694.

Casalou, C., A. Faustino, and D.C. Barral. 2016. Arf proteins in cancer cell migration. Small GTPases, 7:270–282.

Casalou, C., A. Ferreira, and D.C. Barral. 2020. The Role of ARF Family Proteins and Their Regulators and Effectors in Cancer Progression: A Therapeutic Perspective. Front Cell Dev Biol, 8:217.

Chiang, T.S., H.F. Wu, and F.S. Lee. 2017. ADP-ribosylation factor-like 4C binding to filamin-A modulates filopodium formation and cell migration. Molecular biology of the cell, 28:3013–3028.

Choudhury, A., D.L. Marks, K.M. Proctor, G.W. Gould, and R.E. Pagano. 2006. Regulation of caveolar endocytosis by syntaxin 6-dependent delivery of membrane components to the cell surface. Nature cell biology, 8:317–328.

Corre, T., F.J. Arjona, C. Hayward, S. Youhanna, J.H.F. de Baaij, H. Belge, N. Nägele, H. Debaix, M.G. Blanchard, M. Traglia, S.E. Harris, S. Ulivi, R. Rueedi, D. Lamparter, A. Macé, C. Sala, S. Lenarduzzi, B. Ponte, M. Pruijm, D. Ackermann, G. Ehret, D. Baptista, O. Polasek, I. Rudan, T.W. Hurd, N.D. Hastie, V. Vitart, G. Waeber, Z. Kutalik, S. Bergmann, R. Vargas-Poussou, M. Konrad, P. Gasparini, I.J. Deary, J.M. Starr, D. Toniolo, P. Vollenweider, J.G.J. Hoenderop, R.J.M. Bindels, M. Bochud, and O. Devuyst. 2018. Genome-Wide Meta-Analysis Unravels Interactions between Magnesium Homeostasis and Metabolic Phenotypes. Journal of the American Society of Nephrology: JASN, 29:335–348.

D’Souza-Schorey, C., and P. Chavrier. 2006. ARF proteins: roles in membrane traffic and beyond. Nature reviews. Molecular cell biology, 7:347–358.

Delevoye, C., I. Hurbain, D. Tenza, J.B. Sibarita, S. Uzan-Gafsou, H. Ohno, W.J. Geerts, A.J. Verkleij, J. Salamero, M.S. Marks, and G. Raposo. 2009. AP-1 and KIF13A coordinate endosomal sorting and positioning during melanosome biogenesis. The Journal of cell biology, 187:247–264.

Donaldson, J.G., A. Honda, and R. Weigert. 2005. Multiple activities for Arf1 at the Golgi complex. Biochimica et Biophysica Acta (BBA) - Molecular Cell Research, 1744:364–373.

Donaldson, J.G., and C.L. Jackson. 2011. ARF family G proteins and their regulators: roles in membrane transport, development and disease. Nature reviews. Molecular cell biology, 12:362–375.

Donaldson, J.G., and R.D. Klausner. 1994. ARF: a key regulatory switch in membrane traffic and organelle structure. Current opinion in cell biology, 6:527–532.

Dykes, S.S., A.L. Gray, D.T. Coleman, M. Saxena, C.A. Stephens, J.L. Carroll, K. Pruitt, and J.A. Cardelli. 2016. The Arf-like GTPase Arl8b is essential for three-dimensional invasive growth of prostate cancer in vitro and xenograft formation and growth in vivo. Oncotarget, 7:31037–31052.

Edelheit, O., A. Hanukoglu, and I. Hanukoglu. 2009. Simple and efficient site-directed mutagenesis using two single-primer reactions in parallel to generate mutants for protein structure-function studies. BMC Biotechnol, 9:61.

Fisher, S., D. Kuna, T. Caspary, R.A. Kahn, and E. Sztul. 2020. ARF family GTPases with links to cilia. American Journal of Physiology-Cell Physiology, 319:C404–C418.

Francis, J.W., L.E. Newman, L.A. Cunningham, and R.A. Kahn. 2017. A Trimer Consisting of the Tubulin-specific Chaperone D (TBCD), Regulatory GTPase ARL2, and beta-Tubulin Is Required for Maintaining the Microtubule Network. The Journal of biological chemistry, 292:4336–4349.

Fujii, S., S. Matsumoto, S. Nojima, E. Morii, and A. Kikuchi. 2015. Arl4c expression in colorectal and lung cancers promotes tumorigenesis and may represent a novel therapeutic target. Oncogene, 34:4834–4844.

Furman, C., S.M. Short, R.R. Subramanian, B.R. Zetter, and T.M. Roberts. 2002. DEF-1/ASAP1 is a GTPase-activating protein (GAP) for ARF1 that enhances cell motility through a GAP-dependent mechanism. The Journal of biological chemistry, 277:7962–7969.

Gillingham, A.K., and S. Munro. 2007. The small G proteins of the Arf family and their regulators. Annual review of cell and developmental biology, 23:579–611.

Glessner, J.T., J.P. Bradfield, K. Wang, N. Takahashi, H. Zhang, P.M. Sleiman, F.D. Mentch, C.E. Kim, C. Hou, K.A. Thomas, M.L. Garris, S. Deliard, E.C. Frackelton, F.G. Otieno, J. Zhao, R.M. Chiavacci, M. Li, J.D. Buxbaum, R.I. Berkowitz, H. Hakonarson, and S.F.A. Grant. 2010. A Genome-wide Study Reveals Copy Number Variants Exclusive to Childhood Obesity Cases. The American Journal of Human Genetics, 87:661–666.

Hofmann, I., and S. Munro. 2006. An N-terminally acetylated Arf-like GTPase is localised to lysosomes and affects their motility. Journal of cell science, 119:1494–1503.

Howley, B.V., L.A. Link, S. Grelet, M. El-Sabban, and P.H. Howe. 2018. A CREB3-regulated ER-Golgi trafficking signature promotes metastatic progression in breast cancer. Oncogene, 37:1308–1325.

Huang, D., Y. Pei, C. Dai, Y. Huang, H. Chen, X. Chen, X. Zhang, C. Lin, H. Wang, R. Zhang, X. Wan, and L. Wang. 2019. Up-regulated ADP-Ribosylation factor 3 promotes breast cancer cell proliferation through the participation of FOXO1. Experimental cell research, 384:111624.

Jackson, C.L. 2018. Activators and Effectors of the Small G Protein Arf1 in Regulation of Golgi Dynamics During the Cell Division Cycle. Front Cell Dev Biol, 6:29.

Jani, R.A., L.K. Purushothaman, S. Rani, P. Bergam, and S.R. Setty. 2015. STX13 regulates cargo delivery from recycling endosomes during melanosome biogenesis. Journal of cell science, 128:3263–3276.

Jumper, J., R. Evans, A. Pritzel, T. Green, M. Figurnov, O. Ronneberger, K. Tunyasuvunakool, R. Bates, A. Žídek, A. Potapenko, A. Bridgland, C. Meyer, S.A.A. Kohl, A.J. Ballard, A. Cowie, B. Romera-Paredes, S. Nikolov, R. Jain, J. Adler, T. Back, S. Petersen, D. Reiman, E. Clancy, M. Zielinski, M. Steinegger, M. Pacholska, T. Berghammer, S. Bodenstein, D. Silver, O. Vinyals, A.W. Senior, K. Kavukcuoglu, P. Kohli, and D. Hassabis. 2021. Highly accurate protein structure prediction with AlphaFold. Nature, 596:583–589.

Jung, J.J., S.M. Inamdar, A. Tiwari, and A. Choudhury. 2012. Regulation of intracellular membrane trafficking and cell dynamics by syntaxin-6. Bioscience reports, 32:383–391.

Kahn, R.A., J. Cherfils, M. Elias, R.C. Lovering, S. Munro, and A. Schurmann. 2006. Nomenclature for the human Arf family of GTP-binding proteins: ARF, ARL, and SAR proteins. The Journal of cell biology, 172:645–650.

Khatter, D., V.B. Raina, D. Dwivedi, A. Sindhwani, S. Bahl, and M. Sharma. 2015. The small GTPase Arl8b regulates assembly of the mammalian HOPS complex on lysosomes. Journal of cell science, 128:1746–1761.

Klumperman, J., R. Kuliawat, J.M. Griffith, H.J. Geuze, and P. Arvan. 1998. Mannose 6-phosphate receptors are sorted from immature secretory granules via adaptor protein AP-1, clathrin, and syntaxin 6-positive vesicles. The Journal of cell biology, 141:359–371.

Ko, J., H. Park, L. Heo, and C. Seok. 2012. GalaxyWEB server for protein structure prediction and refinement. Nucleic acids research, 40:W294–297.

Krugmann, S., I. Jordens, K. Gevaert, M. Driessens, J. Vandekerckhove, and A. Hall. 2001. Cdc42 induces filopodia by promoting the formation of an IRSp53:Mena complex. Current biology: CB, 11:1645–1655.

Kuliawat, R., J. Klumperman, T. Ludwig, and P. Arvan. 1997. Differential sorting of lysosomal enzymes out of the regulated secretory pathway in pancreatic beta-cells. The Journal of cell biology, 137:595–608.

Kulkarni, A.H., P. Ghosh, A. Seetharaman, P. Kondaiah, and N. Gundiah. 2018. Traction cytometry: regularization in the Fourier approach and comparisons with finite element method. Soft matter, 14:4687–4695.

Kumari, S., and S. Mayor. 2008. ARF1 is directly involved in dynamin-independent endocytosis. Nature cell biology, 10:30–41.

Li, C.C., T.C. Chiang, T.S. Wu, G. Pacheco-Rodriguez, J. Moss, and F.J. Lee. 2007. ARL4D recruits cytohesin-2/ARNO to modulate actin remodeling. Molecular biology of the cell, 18:4420–4437.

Li, Y., K. Ling, and J. Hu. 2012. The emerging role of Arf/Arl small GTPases in cilia and ciliopathies. Journal of cellular biochemistry, 113:2201–2207.

Lippincott-Schwartz, J., N.B. Cole, and J.G. Donaldson. 1998. Building a secretory apparatus: role of ARF1/COPI in Golgi biogenesis and maintenance. Histochem Cell Biol, 109:449–462.

Luchsinger, C., M. Aguilar, P.V. Burgos, P. Ehrenfeld, and G.A. Mardones. 2018. Functional disruption of the Golgi apparatus protein ARF1 sensitizes MDA-MB-231 breast cancer cells to the antitumor drugs Actinomycin D and Vinblastine through ERK and AKT signaling. PloS one, 13:e0195401.

Marwaha, R., D. Dwivedi, and M. Sharma. 2019. Emerging roles of Arf-like GTP-binding proteins: from membrane trafficking to cytoskeleton dynamics and beyond. Proc Indian Natn Sci Acad, 85:189–212.

Matsuba, R., M. Imamura, Y. Tanaka, M. Iwata, H. Hirose, K. Kaku, H. Maegawa, H. Watada, K. Tobe, A. Kashiwagi, R. Kawamori, and S. Maeda. 2016. Replication Study in a Japanese Population of Six Susceptibility Loci for Type 2 Diabetes Originally Identified by a Transethnic Meta-Analysis of Genome-Wide Association Studies. PloS one, 11:e0154093.

Mierke, C.T., P. Kollmannsberger, D.P. Zitterbart, G. Diez, T.M. Koch, S. Marg, W.H. Ziegler, W.H. Goldmann, and B. Fabry. 2010. Vinculin facilitates cell invasion into three-dimensional collagen matrices. The Journal of biological chemistry, 285:13121–13130.

Mora, R., V.L. Bonilha, A. Marmorstein, P.E. Scherer, D. Brown, M.P. Lisanti, and E. Rodriguez-Boulan. 1999. Caveolin-2 localizes to the golgi complex but redistributes to plasma membrane, caveolae, and rafts when co-expressed with caveolin-1. The Journal of biological chemistry, 274:25708–25717.

Nakai, W., Y. Kondo, A. Saitoh, T. Naito, K. Nakayama, and H.W. Shin. 2013. ARF1 and ARF4 regulate recycling endosomal morphology and retrograde transport from endosomes to the Golgi apparatus. Molecular biology of the cell, 24:2570–2581.

Nobes, C.D., and A. Hall. 1995. Rho, rac, and cdc42 GTPases regulate the assembly of multimolecular focal complexes associated with actin stress fibers, lamellipodia, and filopodia. Cell, 81:53–62.

Pandey, A.K., A. Saxena, S.K. Dey, M. Kanjilal, U. Kumar, and B.K. Thelma. 2021. Correlation between an intronic SNP genotype and ARL15 level in rheumatoid arthritis. Journal of genetics. 100.

Pasqualato, S., L. Renault, and J. Cherfils. 2002. Arf, Arl, Arp and Sar proteins: a family of GTP-binding proteins with a structural device for ‘front-back’ communication. EMBO reports, 3:1035–1041.

Patel, M., T.C. Chiang, V. Tran, F.J. Lee, and J.F. Cote. 2011. The Arf family GTPase Arl4A complexes with ELMO proteins to promote actin cytoskeleton remodeling and reveals a versatile Ras-binding domain in the ELMO proteins family. The Journal of biological chemistry, 286:38969–38979.

Pawar, A., J.A. Meier, A. Dasgupta, N. Diwanji, N. Deshpande, K. Saxena, N. Buwa, S. Inchanalkar, M.A. Schwartz, and N. Balasubramanian. 2016. Ral-Arf6 crosstalk regulates Ral dependent exocyst trafficking and anchorage independent growth signalling. Cellular signalling, 28:1225–1236.

Pennauer, M., K. Buczak, C. Prescianotto-Baschong, and M. Spiess. 2022. Shared and specific functions of Arfs 1-5 at the Golgi revealed by systematic knockouts. The Journal of cell biology. 221.

Petrocca, F., D. Iliopoulos, H.R. Qin, M.S. Nicoloso, S. Yendamuri, S.E. Wojcik, M. Shimizu, G. Di Leva, A. Vecchione, F. Trapasso, A.K. Godwin, M. Negrini, G.A. Calin, and C.M. Croce. 2006. Alterations of the tumor suppressor gene ARLTS1 in ovarian cancer. Cancer research, 66:10287–10291.

Presley, J.F., C. Smith, K. Hirschberg, C. Miller, N.B. Cole, K.J. Zaal, and J. Lippincott-Schwartz. 1998. Golgi membrane dynamics. Molecular biology of the cell, 9:1617–1626.

Richards, J.B., D. Waterworth, S. O’Rahilly, M.F. Hivert, R.J. Loos, J.R. Perry, T. Tanaka, N.J. Timpson, R.K. Semple, N. Soranzo, K. Song, N. Rocha, E. Grundberg, J. Dupuis, J.C. Florez, C. Langenberg, I. Prokopenko, R. Saxena, R. Sladek, Y. Aulchenko, D. Evans, G. Waeber, J. Erdmann, M.S. Burnett, N. Sattar, J. Devaney, C. Willenborg, A. Hingorani, J.C. Witteman, P. Vollenweider, B. Glaser, C. Hengstenberg, L. Ferrucci, D. Melzer, K. Stark, J. Deanfield, J. Winogradow, M. Grassl, A.S. Hall, J.M. Egan, J.R. Thompson, S.L. Ricketts, I.R. König, W. Reinhard, S. Grundy, H.E. Wichmann, P. Barter, R. Mahley, Y.A. Kesaniemi, D.J. Rader, M.P. Reilly, S.E. Epstein, A.F. Stewart, C.M. Van Duijn, H. Schunkert, K. Burling, P. Deloukas, T. Pastinen, N.J. Samani, R. McPherson, G. Davey Smith, T.M. Frayling, N.J. Wareham, J.B. Meigs, V. Mooser, and T.D. Spector. 2009. A genome-wide association study reveals variants in ARL15 that influence adiponectin levels. PLoS genetics, 5:e1000768.

Ried, J.S., M.J. Jeff, A.Y. Chu, J.L. Bragg-Gresham, J. van Dongen, J.E. Huffman, T.S. Ahluwalia, G. Cadby, N. Eklund, J. Eriksson, T. Esko, M.F. Feitosa, A. Goel, M. Gorski, C. Hayward, N.L. Heard-Costa, A.U. Jackson, E. Jokinen, S. Kanoni, K. Kristiansson, Z. Kutalik, J. Lahti, J. Luan, R. Mägi, A. Mahajan, M. Mangino, C. Medina-Gomez, K.L. Monda, I.M. Nolte, L. Pérusse, I. Prokopenko, L. Qi, L.M. Rose, E. Salvi, M.T. Smith, H. Snieder, A. Stancáková, Y. Ju Sung, I. Tachmazidou, A. Teumer, G. Thorleifsson, P. van der Harst, R.W. Walker, S.R. Wang, S.H. Wild, S.M. Willems, A. Wong, W. Zhang, E. Albrecht, A. Couto Alves, S.J. Bakker, C. Barlassina, T.M. Bartz, J. Beilby, C. Bellis, R.N. Bergman, S. Bergmann, J. Blangero, M. Blüher, E. Boerwinkle, L.L. Bonnycastle, S.R. Bornstein, M. Bruinenberg, H. Campbell, Y.I. Chen, C.W. Chiang, P.S. Chines, F.S. Collins, F. Cucca, L.A. Cupples, F. D’Avila, E.J. de Geus, G. Dedoussis, M. Dimitriou, A. Döring, J.G. Eriksson, A.E. Farmaki, M. Farrall, T. Ferreira, K. Fischer, N.G. Forouhi, N. Friedrich, A.P. Gjesing, N. Glorioso, M. Graff, H. Grallert, N. Grarup, J. Gräßler, J. Grewal, A. Hamsten,M.N. Harder, C.A. Hartman, M. Hassinen, N. Hastie, A.T. Hattersley, A.S. Havulinna, M. Heliövaara, H. Hillege, A. Hofman, O. Holmen, et al. 2016. A principal component meta-analysis on multiple anthropometric traits identifies novel loci for body shape. Nature communications, 7:13357.

Rocha, N., F. Payne, I. Huang-Doran, A. Sleigh, K. Fawcett, C. Adams, A. Stears, V. Saudek, S. O’Rahilly, I. Barroso, and R.K. Semple. 2017. The metabolic syndrome-associated small G protein ARL15 plays a role in adipocyte differentiation and adiponectin secretion. Scientific reports, 7:17593.

Roy, A., A. Kucukural, and Y. Zhang. 2010. I-TASSER: a unified platform for automated protein structure and function prediction. Nature protocols, 5:725–738.

Sáenz, J.B., W.J. Sun, J.W. Chang, J. Li, B. Bursulaya, N.S. Gray, and D.B. Haslam. 2009. Golgicide A reveals essential roles for GBF1 in Golgi assembly and function. Nature chemical biology, 5:157–165.

Schlienger, S., R.A. Ramirez, and A. Claing. 2015. ARF1 regulates adhesion of MDA-MB-231 invasive breast cancer cells through formation of focal adhesions. Cellular signalling, 27:403–415.

Scott, R.A., T. Fall, D. Pasko, A. Barker, S.J. Sharp, L. Arriola, B. Balkau, A. Barricarte, I. Barroso, H. Boeing, F. Clavel-Chapelon, F.L. Crowe, J.M. Dekker, G. Fagherazzi, E. Ferrannini, N.G. Forouhi, P.W. Franks, D. Gavrila, V. Giedraitis, S. Grioni, L.C. Groop, R. Kaaks, T.J. Key, T. Kuhn, L.A. Lotta, P.M. Nilsson, K. Overvad, D. Palli, S. Panico, J.R. Quiros, O. Rolandsson, N. Roswall, C. Sacerdote, N. Sala, M.J. Sanchez, M.B. Schulze, A. Siddiq, N. Slimani, I. Sluijs, A.M. Spijkerman, A. Tjonneland, R. Tumino, A.D. van der, H. Yaghootkar, R.s. group, E.P.-I. consortium, M.I McCarthy, R.K. Semple, E. Riboli, M. Walker, E. Ingelsson, T.M. Frayling, D.B. Savage, C. Langenberg, and N.J. Wareham. 2014. Common genetic variants highlight the role of insulin resistance and body fat distribution in type 2 diabetes, independent of obesity. Diabetes, 63:4378–4387.

Scott, R.A., V. Lagou, R.P. Welch, E. Wheeler, M.E. Montasser, J. Luan, R. Magi, R.J. Strawbridge, E. Rehnberg, S. Gustafsson, S. Kanoni, L.J. Rasmussen-Torvik, L. Yengo, C. Lecoeur, D. Shungin, S. Sanna, C. Sidore, P.C. Johnson, J.W. Jukema, T. Johnson, A. Mahajan, N. Verweij, G. Thorleifsson, J.J. Hottenga, S. Shah, A.V. Smith, B. Sennblad, C. Gieger, P. Salo, M. Perola, N.J. Timpson, D.M. Evans, B.S. Pourcain, Y. Wu, J.S. Andrews, J. Hui, L.F. Bielak, W. Zhao, M. Horikoshi, P. Navarro, A. Isaacs, J.R. O’Connell, K. Stirrups, V. Vitart, C. Hayward, T. Esko, E. Mihailov, R.M. Fraser, T. Fall, B.F. Voight, S. Raychaudhuri, H. Chen, C.M. Lindgren, A.P. Morris, N.W. Rayner, N. Robertson, D. Rybin, C.T. Liu, J.S. Beckmann, S.M. Willems, P.S. Chines, A.U. Jackson, H.M. Kang, H.M. Stringham, K. Song, T. Tanaka, J.F. Peden, A. Goel, A.A. Hicks, P. An, M. Muller-Nurasyid, A. Franco-Cereceda, L. Folkersen, L. Marullo, H. Jansen, A.J. Oldehinkel, M. Bruinenberg, J.S. Pankow, K.E. North, N.G. Forouhi, R.J. Loos, S. Edkins, T.V. Varga, G. Hallmans, H. Oksa, M. Antonella, R. Nagaraja, S. Trompet, I. Ford, S.J. Bakker, A. Kong, M. Kumari, B. Gigante, C. Herder, P.B. Munroe, M. Caulfield, J. Antti, M. Mangino, K. Small, I. Miljkovic, et al. 2012. Large-scale association analyses identify new loci influencing glycemic traits and provide insight into the underlying biological pathways. Nat Genet, 44:991–1005.

Setty, S.R., T.I. Strochlic, A.H. Tong, C. Boone, and C.G. Burd. 2004. Golgi targeting of ARF-like GTPase Arl3p requires its Nalpha-acetylation and the integral membrane protein Sys1p. Nature cell biology, 6:414–419.

Shakya, S., P. Sharma, A.M. Bhatt, R.A. Jani, C. Delevoye, and S.R. Setty. 2018. Rab22A recruits BLOC-1 and BLOC-2 to promote the biogenesis of recycling endosomes. EMBO reports. 19.

Sharer, J.D., J.F. Shern, H. Van Valkenburgh, D.C. Wallace, and R.A. Kahn. 2002. ARL2 and BART enter mitochondria and bind the adenine nucleotide transporter. Molecular biology of the cell, 13:71–83.

Sharma, A., M. Saini, S. Kundu, and B.K. Thelma. 2020. Computational insight into the three-dimensional structure of ADP ribosylation factor like protein 15, a novel susceptibility gene for rheumatoid arthritis. Journal of biomolecular structure & dynamics:1–16.

Shen, J., M. Liu, J. Xu, B. Sun, H. Xu, and W. Zhang. 2019. ARL15 overexpression attenuates high glucose-induced impairment of insulin signaling and oxidative stress in human umbilical vein endothelial cells. Life sciences, 220:127–135.

Shi, M., H.C. Tie, M. Divyanshu, X. Sun, Y. Zhou, B.K. Boh, L.A. Vardy, and L. Lu. 2022. Arl15 upregulates the TGFbeta family signaling by promoting the assembly of the Smad-complex. Elife. 11.

Singh, V., C. Erady, and N. Balasubramanian. 2018. Cell-matrix adhesion controls Golgi organization and function through Arf1 activation in anchorage-dependent cells. Journal of cell science. 131.

Steffen, A., M. Ladwein, G.A. Dimchev, A. Hein, L. Schwenkmezger, S. Arens, K.I. Ladwein, J. Margit Holleboom, F. Schur, J. Victor Small, J. Schwarz, R. Gerhard, J. Faix, T.E. Stradal, C. Brakebusch, and K. Rottner. 2013. Rac function is crucial for cell migration but is not required for spreading and focal adhesion formation. Journal of cell science, 126:4572–4588.

Sztul, E., P.W. Chen, J.E. Casanova, J. Cherfils, J.B. Dacks, D.G. Lambright, F.S. Lee, P.A. Randazzo, L.C. Santy, A. Schürmann, I. Wilhelmi, M.E. Yohe, and R.A. Kahn. 2019. ARF GTPases and their GEFs and GAPs: concepts and challenges. Molecular biology of the cell, 30:1249–1271.

Te Boekhorst, V., L. Preziosi, and P. Friedl. 2016. Plasticity of Cell Migration In Vivo and In Silico. Annual review of cell and developmental biology, 32:491–526.

Thievessen, I., N. Fakhri, J. Steinwachs, V. Kraus, R.S. McIsaac, L. Gao, B.C. Chen, M.A. Baird, M.W. Davidson, E. Betzig, R. Oldenbourg, C.M. Waterman, and B. Fabry. 2015. Vinculin is required for cell polarization, migration, and extracellular matrix remodeling in 3D collagen. FASEB journal: official publication of the Federation of American Societies for Experimental Biology, 29:4555–4567.

Thomsen, S.K., A. Ceroni, M. van de Bunt, C. Burrows, A. Barrett, R. Scharfmann, D. Ebner, M.I. McCarthy, and A.L. Gloyn. 2016. Systematic Functional Characterization of Candidate Causal Genes for Type 2 Diabetes Risk Variants. Diabetes, 65:3805.

Tse, J.R., and A.J. Engler. 2010. Preparation of hydrogel substrates with tunable mechanical properties. Curr Protoc Cell Biol. Chapter 10:Unit 10 16.

Urano, Y., H. Watanabe, S.R. Murphy, Y. Shibuya, Y. Geng, A.A. Peden, C.C.Y. Chang, and T.Y. Chang. 2008. Transport of LDL-derived cholesterol from the NPC1 compartment to the ER involves the trans-Golgi network and the SNARE protein complex. Proceedings of the National Academy of Sciences, 105:16513–16518.

Volpicelli-Daley, L.A., Y. Li, C.J. Zhang, and R.A. Kahn. 2005. Isoform-selective effects of the depletion of ADP-ribosylation factors 1-5 on membrane traffic. Molecular biology of the cell, 16:4495–4508.

Wu, Y., Y. Bai, D.G. McEwan, L. Bentley, D. Aravani, and R.D. Cox. 2021. Palmitoylated small GTPase ARL15 is translocated within Golgi network during adipogenesis. Biology open. 10.

Yaghootkar, H., R.A. Scott, C.C. White, W. Zhang, E. Speliotes, P.B. Munroe, G.B. Ehret, J.C. Bis, C.S. Fox, M. Walker, I.B. Borecki, J.W. Knowles, L. Yerges-Armstrong, C. Ohlsson, J.R. Perry, J.C. Chambers, J.S. Kooner, N. Franceschini, C. Langenberg, M.F. Hivert, Z. Dastani, J.B. Richards, R.K. Semple, and T.M. Frayling. 2014. Genetic evidence for a normal-weight “metabolically obese” phenotype linking insulin resistance, hypertension, coronary artery disease, and type 2 diabetes. Diabetes, 63:4369–4377.

Yang, J., R. Yan, A. Roy, D. Xu, J. Poisson, and Y. Zhang. 2015. The I-TASSER Suite: protein structure and function prediction. Nature methods, 12:7–8.

Yang, J., and Y. Zhang. 2015. I-TASSER server: new development for protein structure and function predictions. Nucleic acids research, 43:W174–W181.

Yu, C.J., and F.J. Lee. 2017. Multiple activities of Arl1 GTPase in the trans-Golgi network. Journal of cell science, 130:1691–1699.

Zhao, J., M. Wang, W. Deng, D. Zhong, Y. Jiang, Y. Liao, B. Chen, and X. Zhang. 2017. ADP-ribosylation factor-like GTPase 15 enhances insulin-induced AKT phosphorylation in the IR/IRS1/AKT pathway by interacting with ASAP2 and regulating PDPK1 activity. Biochem Biophys Res Commun, 486:865–871.

Zolotarov, Y., C. Ma, I. González-Recio, S. Hardy, G.A.C. Franken, N. Uetani, F. Latta, E. Kostantin, J. Boulais, M.P. Thibault, J.F. Côté, I. Díaz-Moreno, A.D. Quintana, J.G.J. Hoenderop, L.A. Martínez-Cruz, M.L. Tremblay, and J.H.F. de Baaij. 2021. ARL15 modulates magnesium homeostasis through N-glycosylation of CNNMs. Cellular and molecular life sciences: CMLS, 78:5427–5445.

