## Supplementary figures and images for "Golgi localized Arl15 regulates cargo transport, cell adhesion and motility"

### Supplementary Figure 1

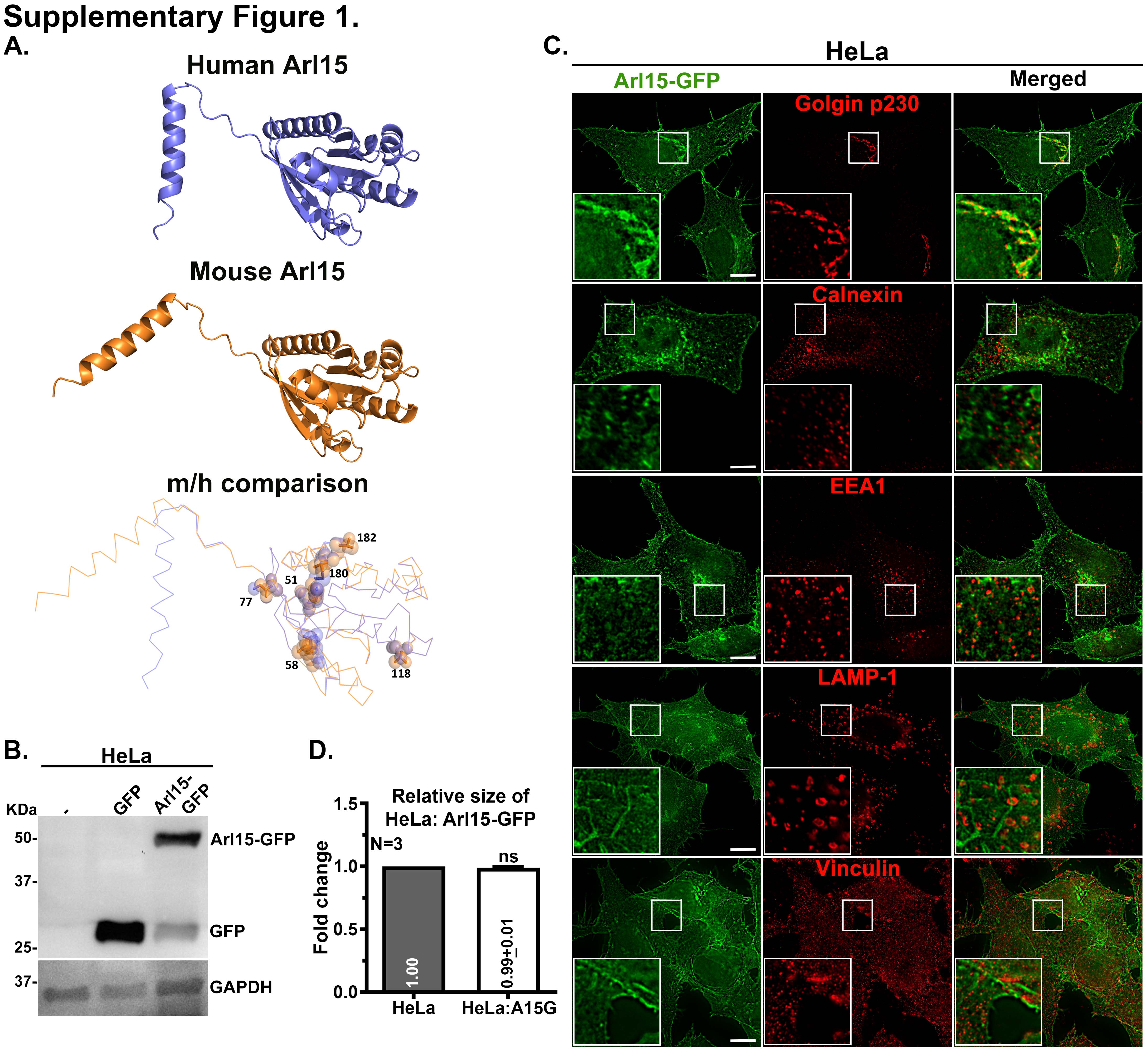

### Supplementary Figure 2

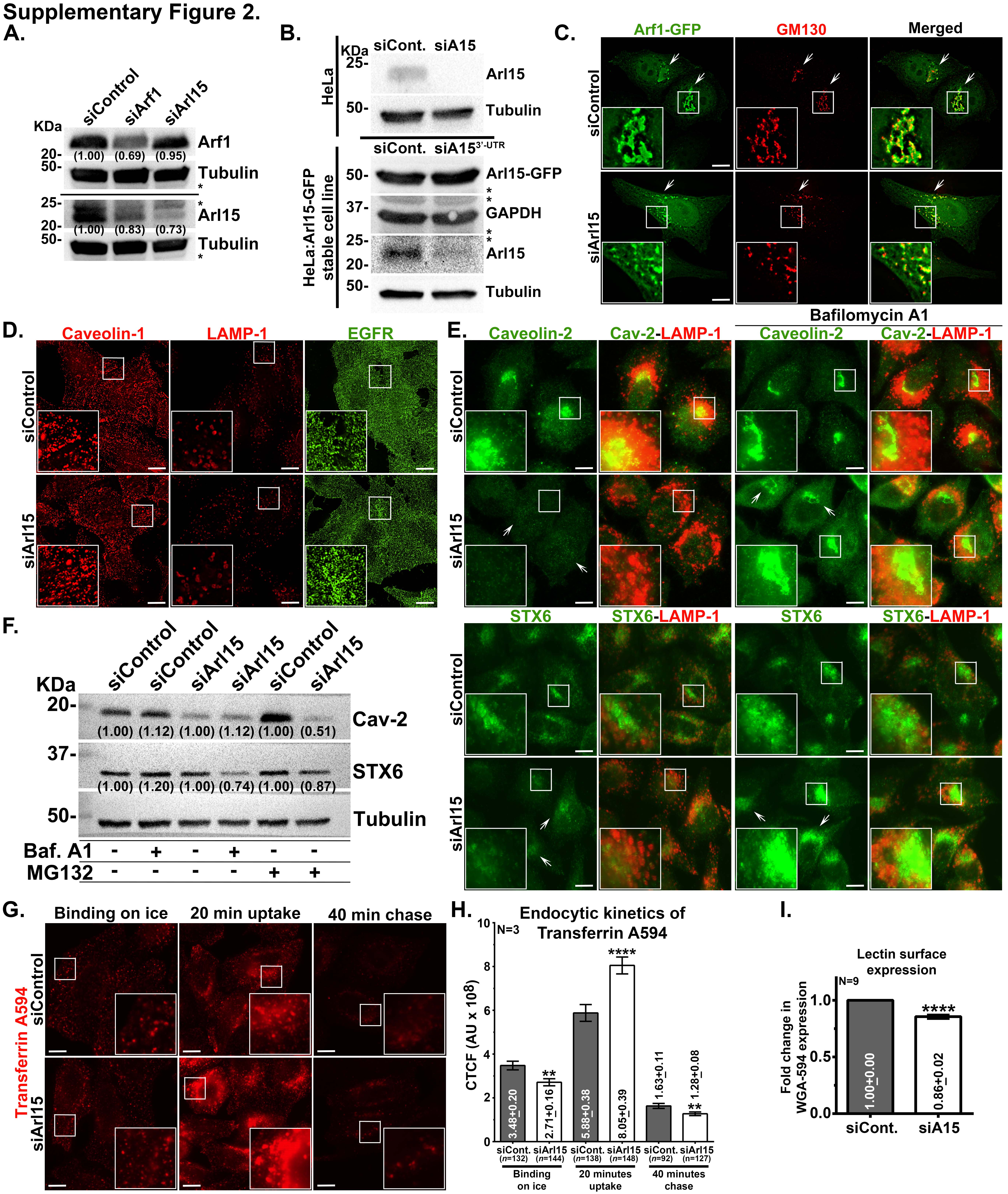

### Supplementary Figure 3

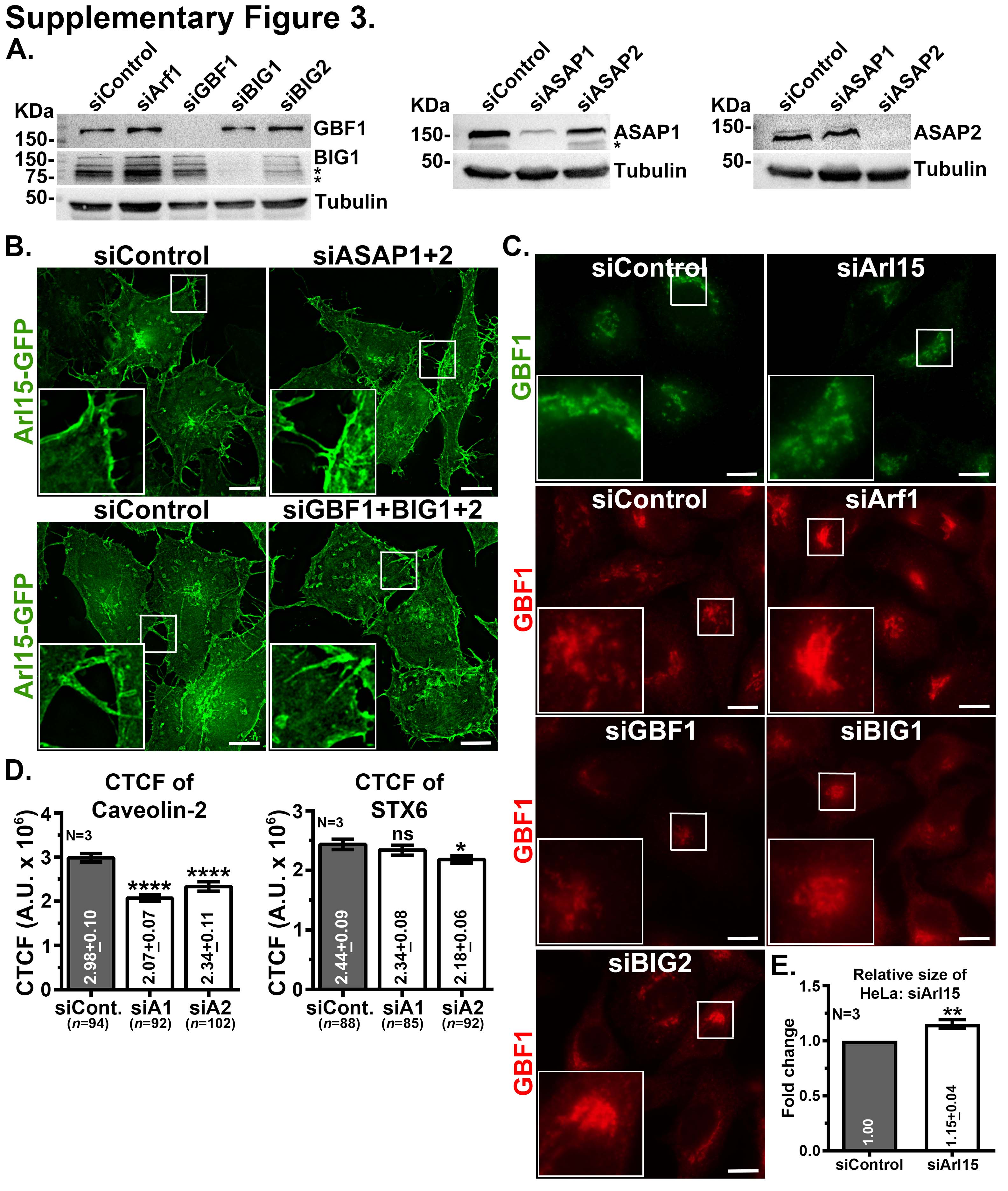
